# USP16 is an ISG15 cross-reactive deubiquitinase targeting a subset of metabolic pathway-related proteins

**DOI:** 10.1101/2023.06.26.546496

**Authors:** Jin Gan, Adán Pinto-Fernández, Dennis Flierman, Jimmy J. L. L. Akkermans, Darragh P. O’Brien, Helene Greenwood, Hannah Claire Scott, Jacques Neefjes, Günter Fritz, Klaus-Peter Knobeloch, Hans van Dam, Benedikt M. Kessler, Huib Ovaa, Paul P. Geurink, Aysegul Sapmaz

## Abstract

The ubiquitin-like modifier ISG15 can modulate host and viral proteins to restrict viral and microbial infections, and act as a cytokine. Its expression and conjugation are strongly up-regulated by type I interferons. Here we identify the deubiquitinating enzyme USP16 as an ISG15 cross-reactive protease. Ubiquitin-specific protease 16 (USP16) was found to react with an ISG15 activity-based probe in pull-down experiments using chronic myeloid leukaemia-derived human cells (HAP1). Supporting this finding, recombinant USP16 cleaved pro-ISG15 and ISG15 iso-peptide linked model substrates *in vitro*, as well as ISGylated substrates present in cell lysates. Moreover, the interferon-induced stimulation of ISGylation in human HAP1 cells was increased by knockdown or knockout of USP16. Depletion of USP16 did not affect interferon signaling, and interferon treatment did not affect USP16 expression or enzymatic activity either. A USP16-dependent ISG15 interactome was established by anti-ISG15 immunoprecipitation mass spectrometry (IP-MS), which indicated that the deISGylating function of USP16 may regulate metabolic pathways involving GOT1, ALDOA, SOD1 and MDH1, all of which were further confirmed to be deISGylated by USP16 in HEK293T cells. Together, our results indicate that USP16 may contribute to regulating the ISGylation status of a subset of proteins related to metabolism during type I interferon responses.

## INTRODUCTION

The innate immune system serves as a first line of defence against viral and bacterial infections in mammalian cells. Activation of this system initiates the release of Type-I interferons (mainly IFN-α and IFN-β), which eventually leads to the transcription of more than 300 interferon-stimulated genes (ISGs). These ISGs encode for different proteins, such as cytokines, chemokines, transcription factors, and enzymes that regulate the host immune response. One of the most strongly induced proteins is ISG15 (interferon-stimulated gene of ∼15 kDa), a ubiquitin(Ub)-like modifier containing 157 amino acids that resembles a linear ubiquitin dimer [1, 2].

ISG15 is translated as 17.8 kDa precursor protein and subsequently processed to liberate its carboxy-terminal LRLRGG motif and become mature ISG15 [3]. Unconjugated mature ISG15 can act as a cytokine [4–7], or interact with intracellular proteins, thereby modulating their functions [8, 9] and stability [10]. Mature ISG15 can also be conjugated to the ε-amine of other proteins via the ISG15 conjugation cycle (ISGylation), a process analogous to ubiquitination. ISG15 conjugation is mediated by a sequential E1-E2-E3 enzyme cascade including E1 UBE1L [11], E2 UBE2L6 [12, 13], and E3s HERC5 [14], TRIM25 [15] and HHARI [16] in human cells, and mHERC6 [17, 18] in murine cells. ISGylation of proteins can regulate their turnover [19–21] and influence complex formation [22, 23].

ISG15 is deconjugated by specific proteases in a process termed deISGylation, comparable to ubiquitin cleavage by deubiquitinating enzymes (DUBs). The major deISGylating enzyme identified in mammalian cells is USP18 [24], a member of the ubiquitin specific protease (USP) family, which cannot cleave ubiquitin, and whose substrate specificity, protein structure, and role in innate immunity have been widely studied [25–28]. USP18 processes pro-ISG15, however, *in vivo* inactivation of USP18 protease activity does not impair the ISGylation machinery [74], suggesting that other proteases with deISGylase activity exist. Since ubiquitin and ISG15 have a common protein fold and an identical C-terminal tail (LRLRGG), it is very likely that additional, Ub-deconjugating, enzymes can process pro-ISG15 into its mature form. Indeed, an initial *in vitro* study [29] and two recent studies [30, 31] identified USP2, USP5, USP13, USP14 and USP21 to react covalently with ISG15-based probes. Among those, deISGylase activity of USP5, USP14 and USP21 was confirmed [29, 31–33]. However, in all studies conducted so far, the cellular function of this ISG15-crossreactivity of certain DUBs still remains elusive. For this reason and because of the observation that pro-ISG15 processing still takes place in the absence of USP18 protease activity, we opted to further explore ISG15-crossreactive DUBs and investigate the biological relevance of their deISGylase activity.

In this study, to identify deISGylating enzymes in an unbiased manner, we used ISG15 activity-based protein profiling (ABPP) assays to detect deISGylating enzymes in human HAP1 cell lysates. USP18, USP5 and USP14 were further identified as deISGylating enzymes in this assay, corroborating previous studies. Interestingly, we identified USP16, a DUB described to regulate the stability of H2A [34], RPS27a [35], and IKKβ [36], as a potential novel ISG15-crossreactive DUB. We found USP16 to be able to cleave pro-ISG15 and isopeptide-linked ISG15-based fluorescence polarization (FP) substrates *in vitro,* as well as natural ISGylated substrates in a cell lysate. To identify ISGylated substrates of USP16 we performed an ISG15 interactome via endogenous ISG15 immunoprecipitation by anti-ISG15 in HAP1 cells lysate combined with mass spectrometry analysis following type I interferon (IFN-I) treatment and using USP16 knockout cells as controls. Malate dehydrogenase, cytoplasmic (MDH1), Superoxide dismutase (SOD) 1, Fructose-bisphosphate aldolase A (ALDOA), and Aspartate aminotransferase, cytoplasmic (GOT1) were identified and further confirmed as ISGylated substrates targeted by USP16, suggesting its potential role as deISGylase in immunometabolic pathways.

## RESULTS

### Activity-based pull-down assay reveals USP16 as an ISG15 cross-reactive DUB

To identify potential ISG15 reactive proteases, we performed a pull-down assay with an ISG15 activity-based probe in lysates from human HAP1 cells with or without interferon-α2 (IFN) stimulation. Proteases reacting with the biotin-tagged human C-terminal domain ISG15 propargylamide (Biotin-hISG15_CTD_-PA) probe [37] were enriched by NeutrAvidin beads, and analysed by label-free LC-MS/MS following washing, elution, and in-solution trypsin digestion (Figure 1A, Data Table S1). Pull-down efficiency was confirmed by immunoblotting using an anti-biotin antibody prior to LC-MS/MS analysis (Supplementary Figure 1). The LC-MS/MS data were processed by cross-comparative analysis of Biotin-hISG15_CTD_-PA versus no-probe samples (Figure 1B), and cross-comparative analysis of IFN-stimulated versus non-stimulated cells (Figure 1C). In the lysates from the IFN-stimulated cells four proteases, USP18, USP14, USP5, and USP16, were significantly enriched in the hISG15_CTD_-PA probe-treated samples as compared to no-probe samples (Figure 1B). USP18 was the only hISG15_CTD_-PA-reacting protease that was significantly enriched in the IFN-stimulated cells as compared to non-stimulated cells (Figure 1C).

**Figure 1.**
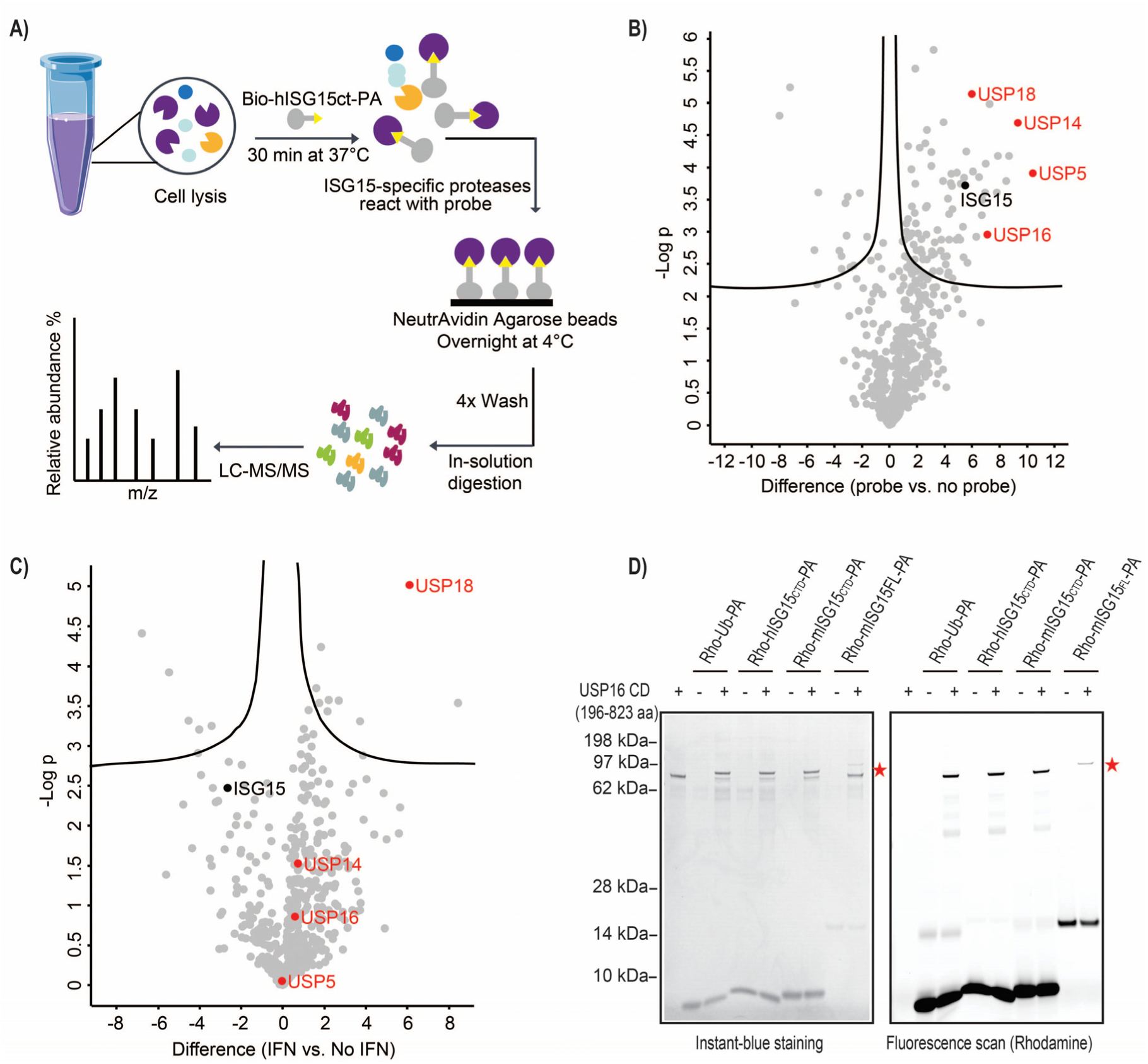
USP16 is pulled-down from human HAP1 cell lysates by Biotin-hISG15_CTD_-PA probe. (A) Streamlined workflow for identification of ISG15-reactive proteases in lysates of HAP1 cells. (B, C) Volcano plots of comparative proteomic analysis of the trypsin digests of the Streptavidin beads pulldowns. Comparison of biotin-based immunoprecipitation of Biotin-hISG15_CTD_-PA probe labeled versus unlabeled IFN-α2 stimulated HAP1 WT cell lysate (B), and comparison of biotin-based immunoprecipitation of IFN-α2 stimulated versus unstimulated cell lysates labeled with Biotin-hISG15_CTD_-PA probe (C). The identified Ubiquitin/ISG15 proteases are shown in red. The statistical cut-off values used for the proteomic analysis are FDR: 0.01 and s0: 0.1. (D) 5μM of recombinant human USP16 catalytic domain (CD, aa 196-823) reacts with 5μM of Rhodamine-tagged Ub-PA, as well as human ISG15_CTD_-PA, mouse ISG15_CTD_-PA and mouse full-length ISG15-PA probes in 10 μL volume at 37 °C for 1 hour. After probe reaction, samples were denatured, resolved on SDS-PAGE, scanned for fluorescence, and stained with InstantBlue Coomassie dye. The probe-labeled USP16 CD is indicated by red stars.

These data were consistent with the previous identification of USP18 as an interferon-inducible deISGylase [38, 39], and the ability of USP18, USP14, and USP5 to react with a human ISG15 full-length activity-based probe [31, 32, 40]. Since USP16 is a so-far unreported ISG15-crossreactive DUB, we validated its ability to bind to both Ub- and ISG15-based probes by incubating purified recombinant human USP16 catalytic domain (CD; aa 196-823) with Ub-PA and ISG15 activity-based probes human ISG15_CTD_-PA [26], mouse ISG15_CTD_-PA [26] and full-length mouse ISG15_FL_-PA [41] probes followed by SDS-PAGE. Binding to Ub and ISG15 based probes was confirmed by the appearance of the corresponding USP16-probe adducts (Figure 1D).

### USP16 cleaves ISG15-related substrates *in vitro*

To determine the potential catalytic activity of USP16 towards ISG15, we tested several *in vitro* substrates that can be recognized and converted by deISGylases. We first investigated whether USP16 can cleave the 1-165 amino acid precursor form of human ISG15 (pro-ISG15) [3, 42]. For this, we used purified recombinant human full-length USP16 (FL; aa 22-823) and human USP16 catalytic domain (CD; aa 196-823). We also included recombinant USP7 as a negative control and the major deISGylase USP18 (canonical deISGylase) and USP5 (previously reported as ISG15 reactive DUB) as positive controls. The two forms of USP16 cleaved pro-ISG15 as efficiently as USP18, while USP5 was less efficient and, as expected, USP7 did not cleave at all (Figure 2A).

**Figure 2.**
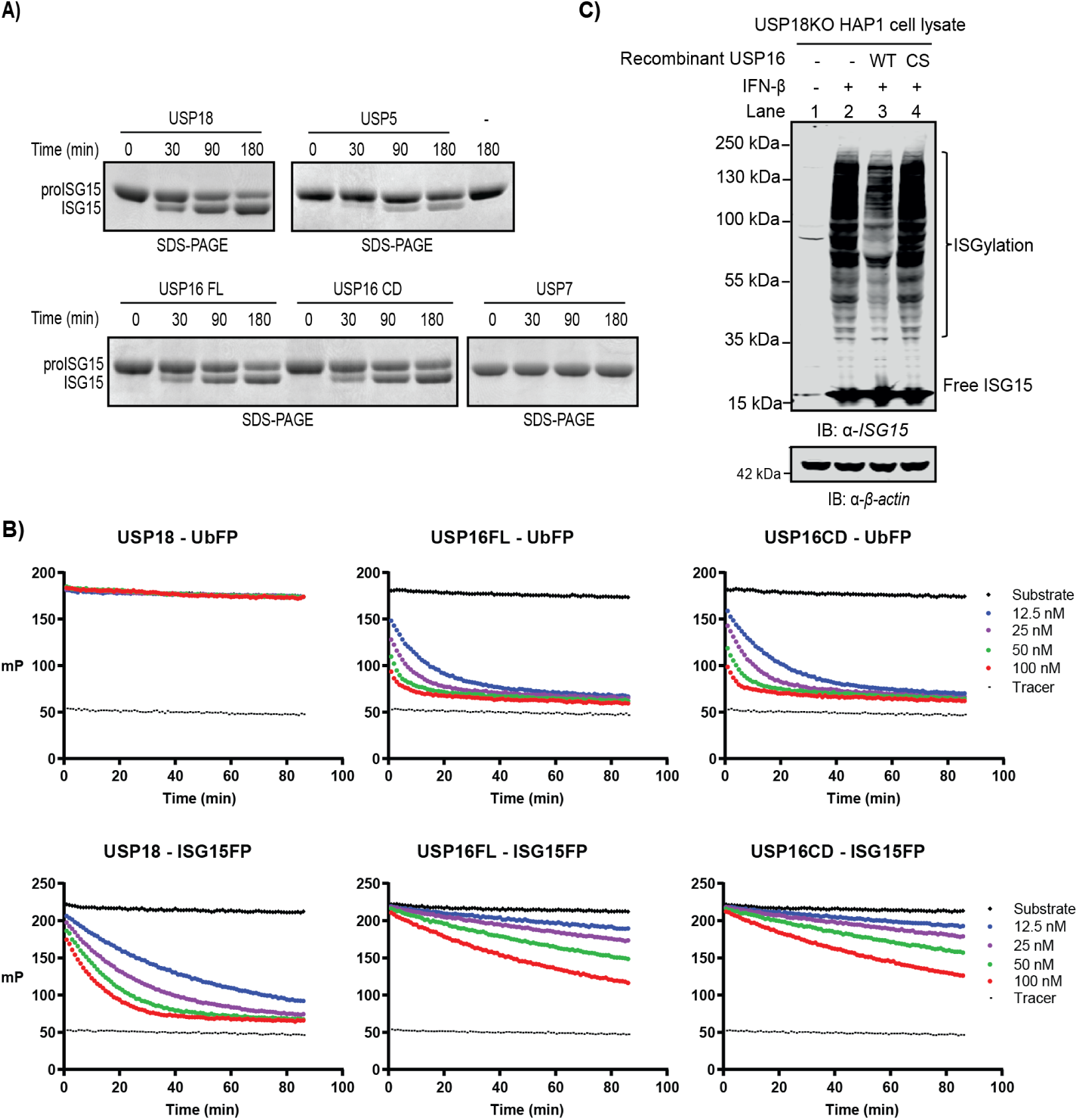
USP16 cleaves ISG15-containing substrates *in vitro*. (A) InstantBlue Coomassie staining of SDS-PAGE of human proISG15 cleaved by recombinant human USP18, USP7, USP5, USP16FL, and USP16CD. (B) The catalytic activity of recombinant human USP18, USP16FL, and USP16CD towards the isopeptide-linked Ub-FP and ISG15-FP substrates. Indicated amounts of USP16 FL/CD enzymes were incubated with 200 nM Ub-FP or ISG15-FP. The substrate cleavage was monitored on the basis of the change in fluorescence polarization (in millipolarization units (mP)). (C) ISG15 deconjugation in lysates of HAP1 USP18KO cells. HAP1 USP18KO cells were stimulated with IFN-β to induce ISGylation on endogenous proteins. 40 μg of cell lysate in 10ul was incubated with recombinant USP16 CD^WT^ or USP16CD^C205S^ at a final concentration of 5uM at 37 °C for 2 hours, and immunoblotted with human ISG15 antibody.

We further assessed the isopeptidase activity of USP16 with Ub- and full-length ISG15-based fluorescence polarization (FP) assay reagents. In these reagents, the C-terminal carboxylate of Ub or full-length ISG15 is linked to the lysine side chain amine of a fluorescent TAMRA-Lys-Gly peptide, thereby mimicking Ub/ISG15 isopeptide-linked substrates [25, 43]. We tested both purified recombinant USP16FL and USP16 CD along with USP18, USP5 and USP7 (Figure 2B, Supplementary Figure 2). USP16 FL and CD started to cleave the ISG15FP substrate at 12.5 nM, while already completely converting the UbFP substrate at this concentration. USP18 processed the ISG15FP substrate more efficiently than USP16 FL and CD did. Notably, USP5 and USP7, which effectively converted the UbFP substrate at subnanomolar concentrations, proved unreactive to the ISG15FP substrate even up to 100 nM concentration. These observations showed that USP16 has clear isopeptidase activity towards ISG15FP substrate *in vitro*.

To determine whether USP16 can remove ISG15 from natural ISGylated substrates, we prepared lysates from interferon-β stimulated HAP1 USP18KO cells (Supplementary Figure 3) and incubated these with purified recombinant USP16CD^WT^ or catalytically inactive mutant USP16CD^C205S^. Wild-type USP16 reduced the amount of ISG15-conjugated proteins in the cell lysate, but as expected, the protease-deficient CS mutant did not, implying that USP16 can effectively process natural ISGylated substrates through its protease function (Figure 2C). These data prove that recombinant USP16 can cleave *in vitro* pro-ISG15, ISG15FP substrate and ISGylated proteins in cell lysates.

### Loss of USP16 leads to elevated accumulation of cellular ISGylation

Having validated USP16 as an ISG15-crossreactive deubiquitinase with *in vitro* substrate cleavage assays, we next investigated the USP16 deISGylation function in living cells. We first examined the effect of siRNA-mediated USP16 depletion. In line with the *in vitro* data, knockdown of USP16 in HAP1 cells with three different siRNAs led to elevated ISGylation after IFN-β stimulation, while the expression level of USP18 remained unchanged (Figure 3A). Next, we used the CRISPR/Cas9 system with guide RNAs targeting either exon 6 (KO #A) or exon 4 (KO #B) of USP16 to generate USP16 knockout (KO) HAP1 cell lines. The knockout was confirmed by USP16-specific Rhodamine-M20-PA probe labeling [44] (Supplementary Figure 4A) and immunoblotting (Supplementary Figure 4B). Like the USP16 siRNA-treated cells, these USP16KO HAP1 cell lines displayed elevated levels of ISGylation after IFN-β stimulation (Figure 3B).

**Figure 3.**
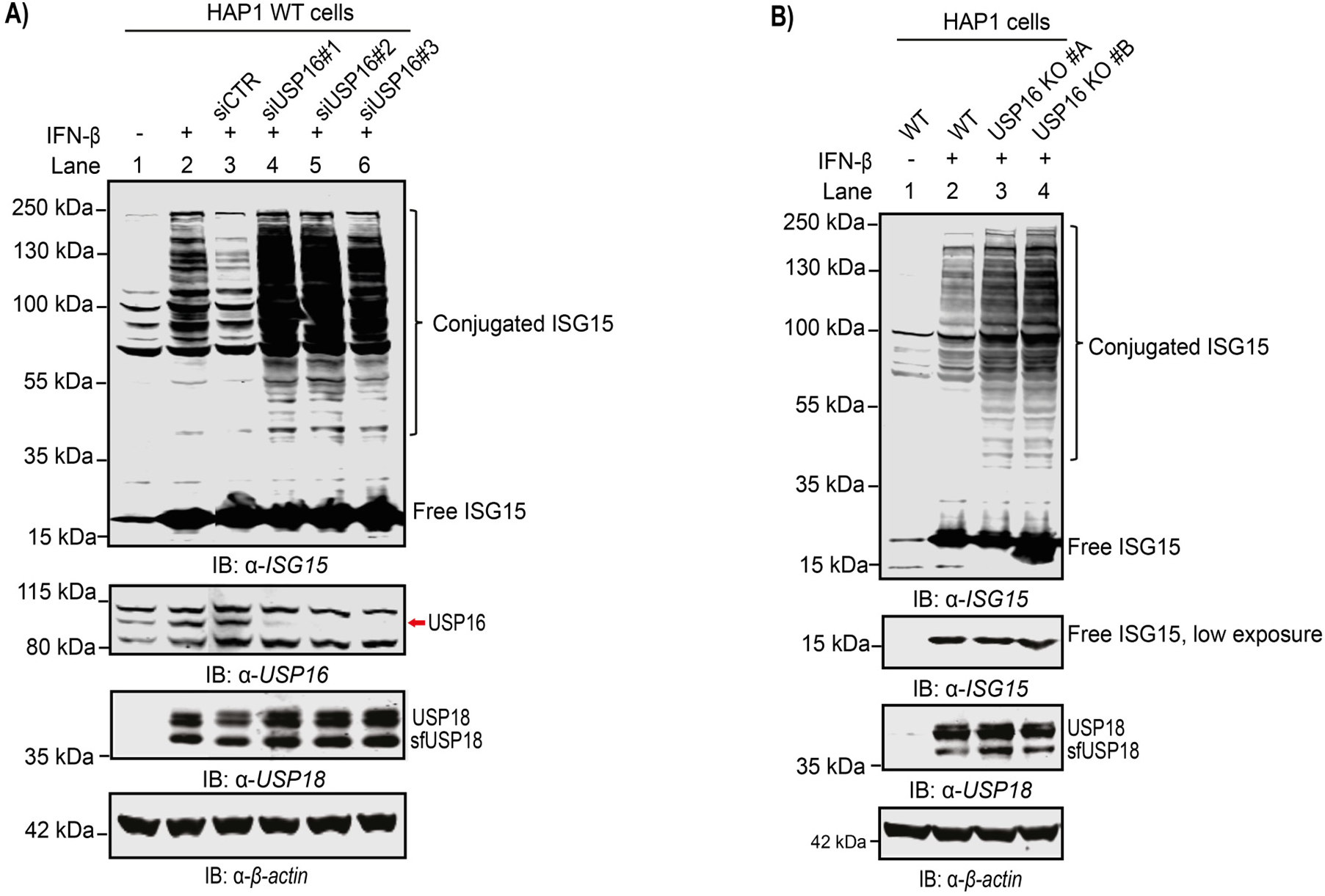
Loss of USP16 leads to elevated cellular ISGylation. (A) Knockdown of USP16. HAP1 WT cells were transfected with either control siRNA (sisiCTR) or three different siRNAs (siUSP16#1, siUSP16#2, siUSP16#3) against USP16 for 72 hours, and 1000 U/ml of IFN-β was added for 24 hours before harvesting. Immunoblot analysis was performed using the indicated antibodies. (B) ISG15 deconjugation in lysates of HAP1 WT and USP16KO cells. Both HAP1 WT and USP16KO cells were stimulated with 1000 U/ml of IFN-β for 24 hours, cell lysates were analysed by immunoblotting using the indicated antibodies.

The observed increased ISGylation in USP16 KO cells could in theory be an indirect effect caused by increased IFN signaling, for instance as a result of a reduced USP18-dependent negative feedback by the USP18-IFNAR2-STAT2 inhibitory complex [45]. Therefore, we assessed whether the USP16 KO HAP1 cells showed increased and/or prolonged IFN-induced phosphorylation of the transcription factors STAT1 and STAT2, or enhanced expression of interferon-stimulated genes [46, 47]. The levels and kinetics of STAT1 and STAT2 phosphorylation did not show clear differences between USP16 KO and WT cells (Supplementary Figure 4C). Moreover, the USP16KO cells did not show changes in interferon-induced ISG15 and USP18 mRNA (Supplementary Figure 4D), nor altered levels of the interferon-induced proteins RIG-I, MDA5, and USP18 (Supplementary Figure 4E). The hISG15ct-PA probe binding activity of cellular deISGlyases was also not affected in the USP16 KO cells (Supplementary Figure 4F). Notably, the protein levels and activity of USP16 were unaffected by IFN-β treatment, in contrast to that of USP18 (Supplementary Figure 5A, B).

Taken together, depletion of USP16 by knockout enhances IFN-β-induced ISGylation, or reduces deISGylation, in the presence of the main ISG15 protease USP18 without affecting interferon signaling. This supports recent findings showing that the deISGlyating functions of USP18, and therefore ISGylation, do not regulate the IFN-I response [48].

### Analysis of the USP16-dependent ISG15 interactome links USP16 with cellular metabolism

To obtain global insights in the cellular pathways and functions regulated by the deISGylase activity of USP16, we established and analysed the endogenous ISG15 interactome in HAP1 WT and HAP1 USP16KO cells using an ISG15 immuno-precipitation mass spectrometry (IP-MS) approach [37]. Duplicate dishes were stimulated with IFN-β, and unstimulated cells were taken along as ISGylation-negative controls. The cell lysates in each sample group were immunoprecipitated with human ISG15 antibody, digested by trypsin, and subjected to label-free quantitative proteomics analysis (Figure 4A). Overall, we identified 2829 proteins (Data Table S2). The ISG15 interactome in HAP1 WT cells (Supplementary Figure 6A) and USP16KO cells (Supplementary Figure 6B) were enriched at least twofold upon IFN-β treatment. Comparison of the ISG15 interactomes of USP16KO and WT cells unveiled a USP16-regulated ISG15 interactome of 142 proteins with at least 1.5 fold changes (Figure 4B, Data Table S3).

**Figure 4.**
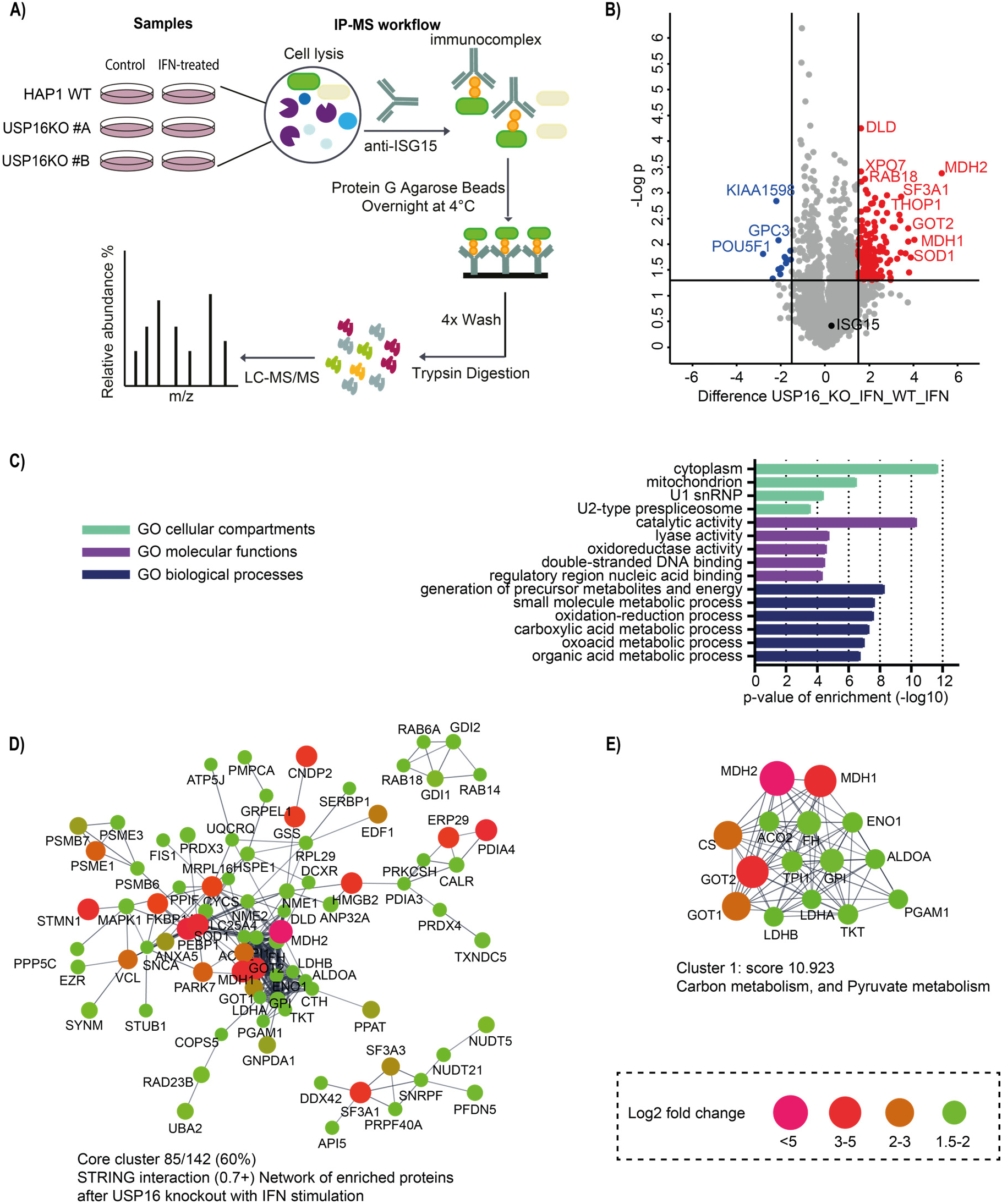
Analysis of the USP16-dependent ISG15 interactome links USP16 with cellular metabolism. (A) Schematic representation of the ISG15 immunoprecipitation mass spectrometry (IP-MS) workflow for analysis of the ISG15 interactome. HAP1 WT and USP16KO cells were stimulated by 1000 U/ml of IFN-β for 48 hours to produce ISGylated substrates as two biological replicates. (B) Volcano plot showing all identified proteins within the IFN-β stimulated USP16KO samples compared with the IFN-β stimulated WT. Dashed lines indicate a cutoff at twofold change (log2 = 1) and a p-value of 0.05 (−log10 = 1.3), n = 2 independent experiments. Source data are provided as Data Table S6. In red are shown the upregulated proteins in the USP16KO cells, named as “USP16-dependent ISG15 interactome”. In blue are shown the downregulated proteins in USP16KO cells. (C) Gene Ontology (GO) enrichment analysis of the USP16-dependent ISG15 interactome. The bar graph shows the most significantly overrepresented GO terms for cellular compartments (CC) in light green, molecular functions (MF) in purple, and biological processes (BP) in dark blue compared against the annotated human proteome. The full GO terms for molecular functions (MF) refer to Supplementary Figure 7A and biological processes (BP) refer to Supplementary Figure 7B. (D) STRING network analysis of the USP16-dependent ISG15 interactome, with a STRING interaction confidence of 0.7 or higher. Cytoscape software was used to visualize the interaction network. Color and node size indicates the fold-change differences in abundance after USP16KO compared with the WT control under IFN-β treatment. (E) Cluster 1 contains multiple proteins involved in carbon metabolism and pyruvate metabolism. MCODE was used to extract the most highly interconnected cluster (cluster 1) from the network shown in (D).

To identify enriched Gene Ontology (GO) terms we performed PANTHER GO-slim analysis, focusing on cellular compartments, molecular functions, and biological processes [49–51]. The analysis confirmed that the identified proteins were strongly enriched for enzymes functioning in metabolic processes and located in cytoplasm and mitochondria (Figure 4C, Supplementary Figure 7). STRING interaction network analysis [52] also revealed a large interconnected set of cytoplasmatic and mitochondrial proteins (Figure 4D). The most interconnected cluster, revealed by the Cytoscape plug-in MCODE [53], consisted of 15 proteins that are involved in carbon metabolism and pyruvate metabolism (Figure 4E). Importantly, all these proteins have been found to be ISGylated in previous proteomic studies [54–60] (Data Table S4). Collectively, our analysis suggests that the USP16-dependent ISG15 interactome is associated with cellular metabolism.

### DeISGylation of discrete substrates by USP16

To examine whether selected members of the identified ISG15 interactome are indeed ISGylated and cleaved by USP16 in cells, we performed in-cell ISGylation assays by co-transfecting hUBE1L, UBCH8, HERC5, and Flag-hISG15 in HEK293T cells [14, 61]. Ectopic expression of USP16 WT, but of not catalytically inactive mutant USP16^C205S^, decreased the overall ISGylation slightly, while expression of USP18 WT had stronger effects (Figure 5A).

**Figure 5.**
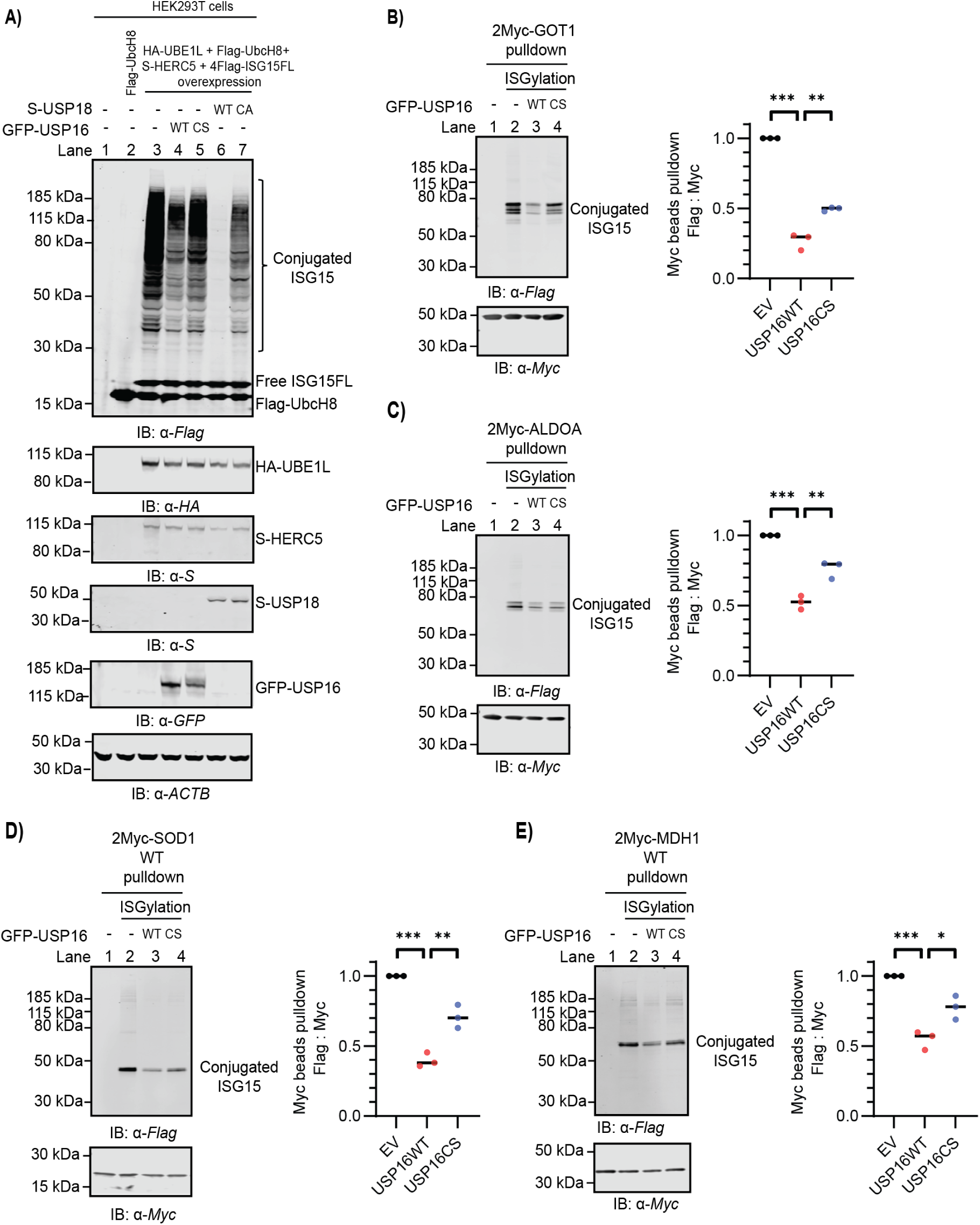
Validation of selected proteins as ISGylated substrates for USP16 enzymatic cleavage. (A) DeISGylation assay upon USP16 or USP18 overexpression in HEK293T cells. Cellular ISGylation was achieved by co-transfecting an ISGylation plasmid mixture including 2 μg HA-UBE1L, 2 μg Flag-UBCH8, 2 μg S-HERC5 and 2 μg HA-ISG15 in HEK293T cells in 6cm dishes for 24 hours. 4 μg Empty vector (in lane 3), or USP16 WT / C205S, or USP18 WT / C64A were co-transfected with the ISGylation plasmids as indicated. Cell lysates were analysed by immunoblotting using the indicated antibodies. (B, C, D, E) Validation of 2Myc-GOT1 (B), 2Myc-ALDOA1 (C), 2Myc-SOD1 (D), and 2Myc-GOT1 (E) as ISGylated substrates for USP16 cleavage. 2 μg 2Myc-tagged protein plasmids were co-transfected with ISGylation plasmid mixture, and the indicated empty vector, GFP-USP16 WT/C205S (amount the same as in 5A) in HEK293T cells in 6cm dishes for 24 hours. Myc-tagged proteins were immunoprecipitated from the cell lysates by Myc trap beads. Precipitated proteins were immunoblotted using the indicated antibodies. Quantification of the ISGylated protein amount in lane 2-4. The ratio of main anti-flag vs. anti-Myc protein band intensity was normalized to empty vector (EV) control. Bar graphs report mean, error bars reflect ±s.d. All significant values were calculated using Student’s t test: **p < 0.05, ***p < 0.001, NS = not significant.

Next, we co-transfected Myc-tagged substrates together with the ISGylation constructs. Following transfection for 24 hours, the cells were lysed and subjected to immunoprecipitation by Myc trap beads, and precipitates were analysed by immunoblotting. We calculated the relative ISGylated protein amounts in these experiments by determining the ratio of the intensity of the ISGylated bands (based on the anti-Flag immunoblot signal) and the intensity of total protein bands (based on the anti-Myc immunoblot signal) and compared the normalized ISGylated protein amounts obtained from the transfected constructs. This indicated that expression of USP16 WT can significantly decrease ISGylation level of GOT1, ALDOA, SOD1 and MDH1. Although expression of USP16^C205S^ affected the ISGylation level to some degree, this was never as significant as USP16 WT (Figure 5B, C, D, E). In summary, GOT1, ALDOA, SOD1 and MDH1 can be ISGylated by the UBE1L-UBCH8-HERC5 cascade in HEK293T cells and are subjected to USP16-dependent cleavage.

## DISCUSSION

Activity-based probe profiling of ubiquitin and ubiquitin-like processing enzymes has become a powerful strategy for monitoring their enzymatic activity changes and identifying novel Ub/Ubl-processing enzymes [62–64] [65]. Moreover, probes generated for different Ubls, led to the discovery of several DUBs that show affinity/activity towards multiple Ubls, which revealing new questions on the biological importance of these cross reactivities. For instance, using ISG15-based ABPP, recombinant USP2, USP5, USP13, USP14 were shown to react covalently with an ISG15-vinylsulfone (ISG15-VS) activity-based probe [29], suggesting that, next to USP18, additional cellular deISG15ylases may exist. Also, USP21 harbours deISGylase activity by processing the ISG15-AMC substrate and deconjugating ISGylated proteins in a lysate derived from IFN-β stimulated HeLa cells [33]. Furthermore, pull-down experiments with immobilized ISG15-propargylamide (ISG15-PA) and ISG15-dehydroalanine (ISG15-Dha) probes resulted in the identification of the previously reported cross-reactive DUBs USP5, USP14 and USP21 [31, 32]. DeISGylase activity for USP5 and USP14 was confirmed using the ISG15-AMC assay. Our study expanded this by identifying USP16 as an ISG15-crossreactive DUB in human HAP1 cell lysates and demonstrating deISGylase activity of USP16 on several substrates subjected to ISGylation with potential links to immunometabolism.

We showed that recombinant USP18, USP16 and USP5 can cleave pro-ISG15. Proteolytic processing of ISG15 precursor protein is necessary to reveal C-terminal GlyGly required for protein conjugation [3, 42]. Since loss of USP18 or USP16 does not reduce ISGylation in cells (Figure 3A, C; Supplementary Figure 3), pro-ISG15 processing is not dependent on a single enzyme. ISG15-crossreactive DUBs, including USP16 and USP5, are likely to compensate for the loss of USP18 to ensure pro-ISG15 cleavage.

Besides performing *in vitro* substrate cleavage assays, further evidence for a deISGylating function of endogenous USP16 in cells was provided by enhanced interferon-stimulated ISGylation in HAP1 cells depleted of USP16 by either knockdown with three different siRNAs or knockout via two different gRNAs. Interferon signaling itself was unaltered in USP16KO HAP1 cells. IFN-β treatment did not affect USP16 protein expression or probe-labeled enzymatic activity in either HeLa and HAP1 cells, indicating that USP16 is not an interferon-inducible gene, in contrast to the main deISGlyase USP18. Moreover, the loss of USP16 did not affect type I interferon signaling, indicating that, unlike USP18 [45], USP16 is not involved in an interferon feed-back mechanism. This also aligns with recent discoveries demonstrating uncloupling of ISGylation from the IFN-I response pathway activation [48].

What is the molecular basis for USP16 cross reactivity? USP16 contains both a Zinc-finger (ZnF) domain (aa 22-143) and a ubiquitin specific protease (USP) domain (aa 196-822). The USP domain is responsible for the catalytic activity of USP16, and the ZnF domain has been reported to recognize free glycine at the C-terminal tail of free ubiquitin, and may function as a sensor for free ubiquitin in cells to regulate ubiquitin-dependent processes [66, 67]. Recombinant human USP16 FL (aa 22-823) and CD (aa 196-823) were found to exhibit similar enzymatic activity towards full-length pro-ISG15 and ISG15-FP. This suggests that the deISGylating activity of recombinant human USP16 is independent of the ZnF domain. However, the ISG15-based reagents used in this study do not contain the free C-terminal tail of ISG15. It is therefore still unclear whether the ZnF domain of USP16 binds free ISG15 in the same mode as free ubiquitin, and if it plays a regulatory role in ISG15-dependent processes in cells.

The specificity of mouse USP18 toward mouse ISG15 is mediated by the interaction between a hydrophobic patch in USP18 and a hydrophobic region in the C-terminal domain of ISG15. The C-terminal domain of ISG15 is necessary and sufficient for USP18 binding [26]. In our study, the C-terminal domain of human ISG15 was sufficient for reaction with human USP5, USP14, USP16, and USP18, indicating that this domain determines ISG15-crossreactivity of DUBs. Interestingly, similar to human USP18 reacting with mouse ISG15-PA and vice versa [26], recombinant human USP16 also displayed cross-species reactivity towards mouse ISG15-PA probe (Figure 1D). However, a high-resolution structure of USP16 in complex with ISG15 is needed to explain the molecular basis for these interactions.

USP16 was initially identified as a DUB that removes ubiquitin from lysine 119 of H2A [34, 68]. It appears to localize primarily in the cytoplasm in cultured human cells [35, 69], and several cytoplasmatic proteins were identified as targets for USP16-mediated deubiquitination, including RPS27a [35], IKKβ [36], and calcineurin A [70]. In line with this, the USP16-dependent ISG15 interactome identified in our experiments mainly contained cytoplasmic proteins, suggesting that USP16 also exerts its deISGylating function primarily in the cytoplasm. Interestingly, none of the reported substrates of USP16-mediated deubiquitination were found in the USP16-dependent ISG15 interactome from IFN-β stimulated HAP1 cells. ISGylation is a key element of the innate immune response by modifying host or viral proteins to restrict the replication or spreading of pathogens [71]. However, emerging evidence showed that ISG15 conjugation is also linked to mitochondrial functions and cellular metabolism [72, 73]. The enrichment of the USP16-dependent ISG15 interactome for metabolic enzymes functioning in cytoplasm and mitochondria also indicates that ISG15 may regulate cellular metabolism and mitochondrial function. This characteristic of the USP16-dependent ISG15-interactome is distinct from the previously studied USP18-dependent ISG15 interactome, in which proteins involved in interferon signaling, innate immune responses and virus defense responses are primarily enriched [37]. Metabolic enzymes GOT1, ALDOA, SOD1 and MDH1 were confirmed here as bona fide ISGylated substrates for USP16 cleavage (Figure 6). Three of the enzymes are involved in gluconeogenesis (GOT1, MDH1 and ALDOA), potentially suggesting functional consequences of USP16-dependent deISGylation events in immunometabolism in general and cellular gluconeogenesis in particular. USP16 antagonists could, therefore, have potential as therapeutic targets for immunometabolic diseases.

**Figure 6.**
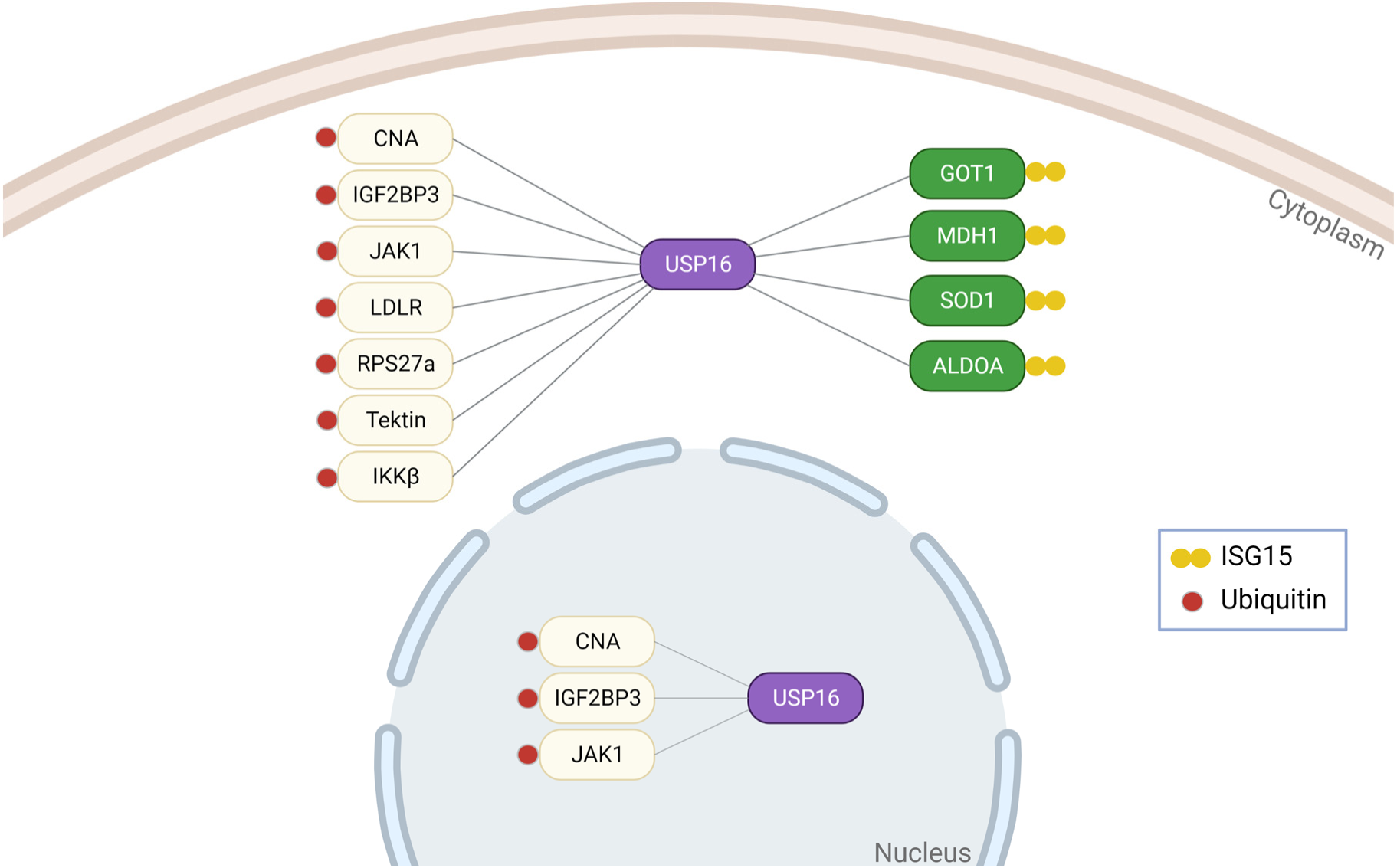
Schematic overview of USP16 roles as dual deubiquitylating and deISGylating enzyme of cellular substrates. Canonical, ubiquitylated protein substrates are colored in yellow, and novel ISGylated metabolic enzyme substrates in green.

## MATERIALS AND METHODS

### Cell culture and transfections

Human HEK293T (Cat# ATCC® CRL-3216™) and HeLa (Cat# ATCC® CCL-2™) cells were cultured under standard conditions in Dulbecco’s modified Eagle’s medium (DMEM) (Gibco) supplemented with 8% FCS (Biowest) and 1% penicillin/streptomycin at 37 °C and 5% CO_2_. Chronic myelogenous leukemia (CML)-derived HAP1 WT (Horizon #C631), HAP1 USP18KO (Horizon #HZGHC000492C011) [1] and HAP1 USP16KO cells (self-made, generated from HAP1 WT) were cultured in Iscove’s Modified Dulbecco’s Medium (IMDM) (Gibco) supplemented with 8% FCS (Biowest), 1% penicillin/streptomycin at 37 °C and 5% CO_2_. All cell lines were tested for mycoplasma contamination using MycoAlert^TM^ Mycoplasma Detection Kit (Lonza, Catalog #: LT07-318) on a monthly basis.

For siRNA transfections, non-targeting siRNA control pools (#D-001206-13-05) and USP16 siRNA oligos (#MQ-006067-01-0002) were purchased from Dharmacon, including siUSP16#1 (Cat# D-006067-01), siUSP16#2 (Cat# D-006067-02), siUSP16#3 (Cat# D-006067-03). Silencing was performed in HAP1 WT cells as follows: for 6-well plate format, 200 µL siRNA (500 nM stock) were incubated with 4 µL Dharmafect reagent 1 (Dharmacon) diluted in 200 µL medium without supplements (total volume of 200 µL transfection mix) with gentle shaking for 20 min at room temperature (RT). A total of 80,000 cells resuspended in 1.6 mL of growth medium were added to transfection mixes to a total volume of 2 mL per well and cultured for 3 days prior to further analysis.

For DNA transfections, HEK293T cells were seeded to achieve 50–60% confluence the following day and transfected using PEI (polyethylenimine, Polysciences Inc., Cat# 23966) as follows: 200 μL DMEM medium without supplements was mixed with DNA and PEI (1 mg/mL) with a ratio at 1:3 (eg: 1μg DNA : 3μL PEI), incubated at RT for 20 min, and added drop-wise to the cells for culturing for 24 h prior to further analysis. Overexpression constructs are listed in Material table 2.

**Material Table 1:**
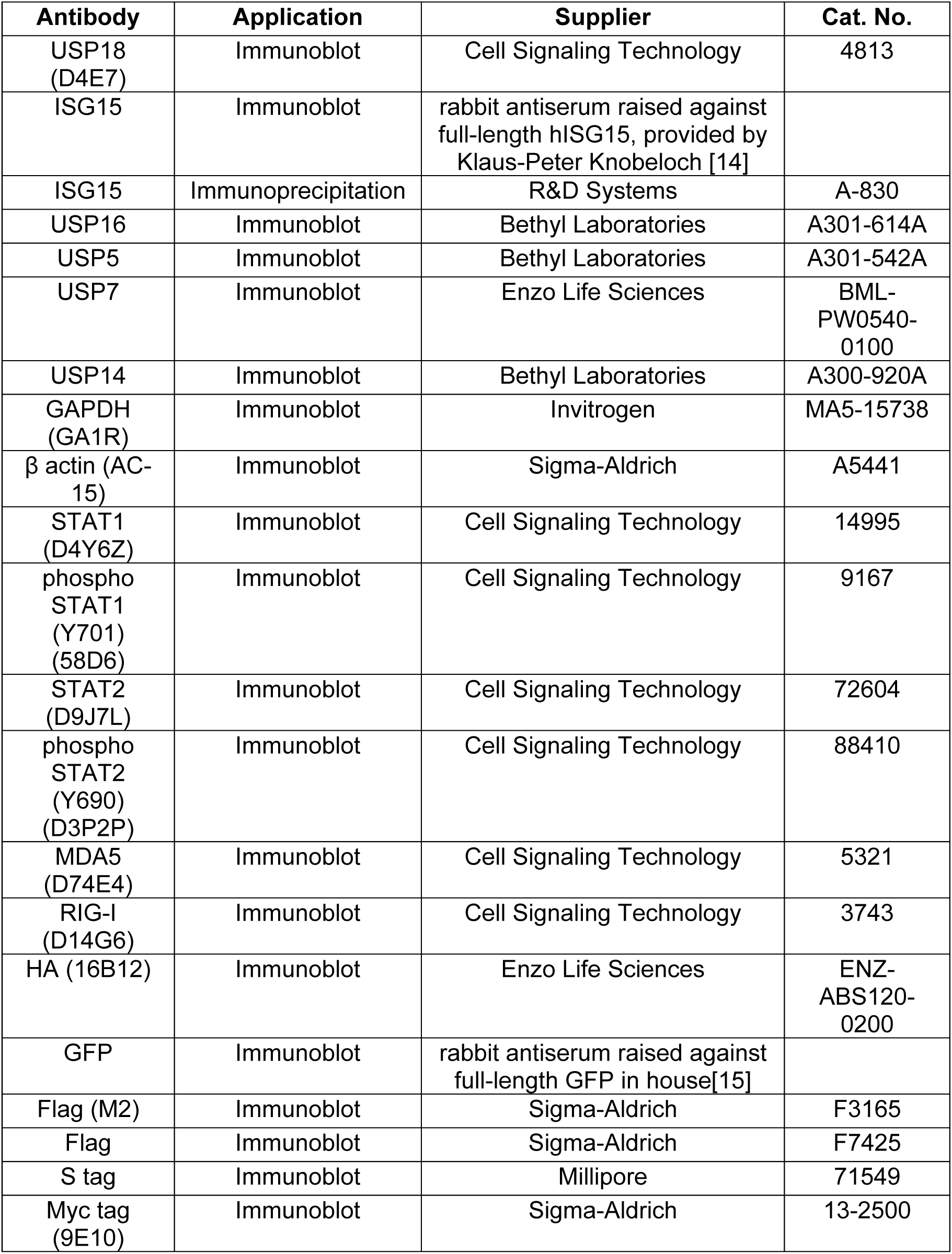
Details of the antibodies utilised in this study.

| **Antibody** | **Application** | **Supplier** | **Cat. No.** |
| --- | --- | --- | --- |
| USP18 (D4E7) | Immunoblot | Cell Signaling Technology | 4813 |
| ISG15 | Immunoblot | rabbit antiserum raised against full-length hISG15, provided by Klaus-Peter Knobeloch [14] |  |
| ISG15 | Immunoprecipitation | R&D Systems | A-830 |
| USP16 | Immunoblot | Bethyl Laboratories | A301-614A |
| USP5 | Immunoblot | Bethyl Laboratories | A301-542A |
| USP7 | Immunoblot | Enzo Life Sciences | BML-PW0540-0100 |
| USP14 | Immunoblot | Bethyl Laboratories | A300-920A |
| GAPDH (GA1R) | Immunoblot | Invitrogen | MA5-15738 |
| $\beta$ actin (AC-15) | Immunoblot | Sigma-Aldrich | A5441 |
| STAT1 (D4Y6Z) | Immunoblot | Cell Signaling Technology | 14995 |
| phospho STAT1 (Y701) (58D6) | Immunoblot | Cell Signaling Technology | 9167 |
| STAT2 (D9J7L) | Immunoblot | Cell Signaling Technology | 72604 |
| phospho STAT2 (Y690) (D3P2P) | Immunoblot | Cell Signaling Technology | 88410 |
| MDA5 (D74E4) | Immunoblot | Cell Signaling Technology | 5321 |
| RIG-I (D14G6) | Immunoblot | Cell Signaling Technology | 3743 |
| HA (16B12) | Immunoblot | Enzo Life Sciences | ENZ-ABS120-0200 |
| GFP | Immunoblot | rabbit antiserum raised against full-length GFP in house[15] |  |
| Flag (M2) | Immunoblot | Sigma-Aldrich | F3165 |
| Flag | Immunoblot | Sigma-Aldrich | F7425 |
| S tag | Immunoblot | Millipore | 71549 |
| Myc tag (9E10) | Immunoblot | Sigma-Aldrich | 13-2500 |

**Material Table 2:**
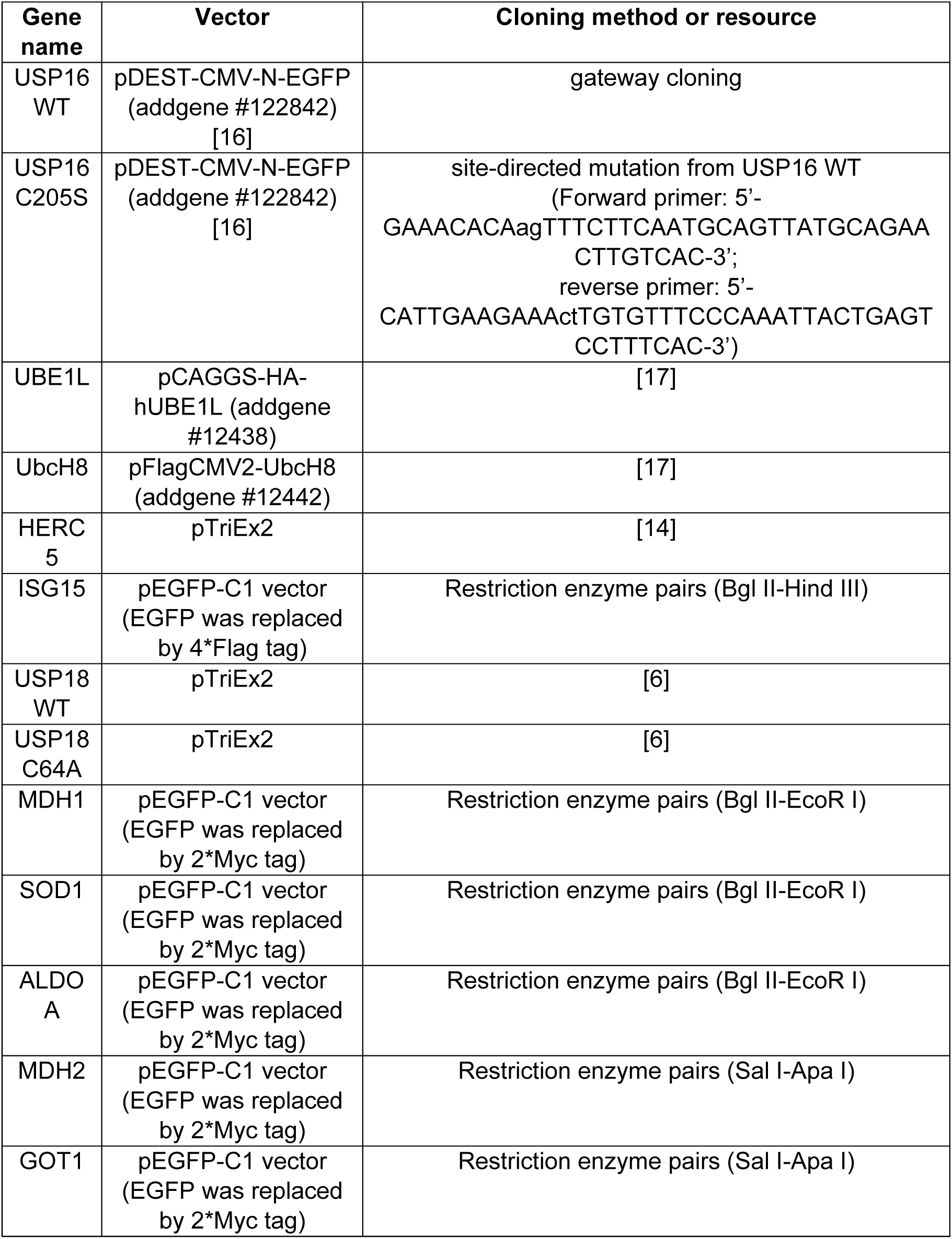
Details of the plasmids utilised in this study.

### Stimulation of cells

Cells were treated with 1000 U/mL of recombinant human IFN-β (PeproTech #300-02BC) or recombinant human IFN-α2 (PBL Assay Science #11105-1) for the indicated times.

### Generation of HAP1 USP16KO cells

HAP1 USP16 KO cell lines were generated using the CRISPR/Cas9 system. The guide RNA sequences targeting exon 4 (USP16 gRNA #B: 5’-CACCGTATTGTCAGTCTTACAGTCT-3’ and 5’-AAACAGACTGTAAGACTGACAATAC) or exon 6 (USP16 gRNA #A: 5’-CACCGAATCAACCACTTGACCCAAC-3’ and 5’-AAACGTTGGGTCAAGTGGTTGATTC-3’) were used as before [2]. Annealed gRNAs were ligated into the lentiCRISPR_v2 vector [3]. HAP1 WT cells were transfected with gRNA-annealed lentiCRISPR_v2 vectors using PEI reagent and selected with puromycin at 1 μg/mL for 3 days. Individual clones were expanded and screened for mutations in the USP16 gene by PCR and immunoblotting. PCR products were sequenced and analysed for indel mutations using ICE CRISPR Analysis Tool from Synthego (https://ice.synthego.com/#/) [4] and by manual inspection of the sequencing profiles with Snapgene.

### Real-time Quantitative RT-PCR

The mRNA level of endogenous USP18 and ISG15 was assessed by RT-qPCR as before [5]. In brief, total RNAs were extracted using the NucleoSpin RNA II kit (MACHEREY-NAGEL) following to the manufacturer instructions. 1ug of total RNA were reversely transcribed using RevertAid First Stand cDNA synthesis Kits (Thermo Fisher), and real-time quantitative PCR experiments were performed using SYBR Green (Promega) in CFX connect Real-Time PCR detection system (Bio-Rad). All the values for target gene expression were normalized to GAPDH. All the experiments were repeated in at least n=3 independent experiments. Primers are listed below:

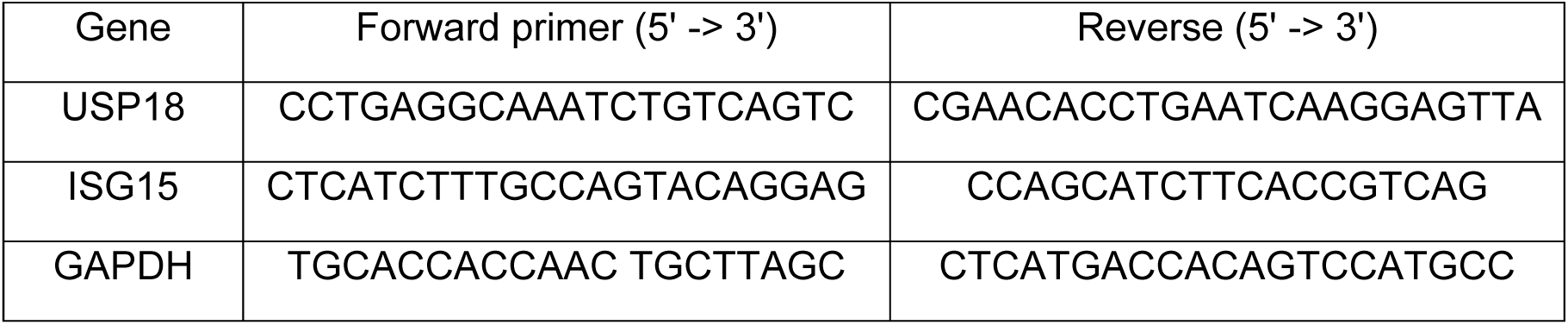

### ISG15-PA, Ub-PA and M20-PA probe labeling assays

The Biotin / Rhodamine-tagged human C terminal domain hISG15_CTD_-PA, Rhodamine-tagged mouse C terminal domain mISG15_CTD_-PA, Biotin-tagged mouse full-length ISG15-PA, Rhodamine-tagged mouse full-length ISG15-PA, Ub-PA, and Rhodamine-tagged M20-PA probe came from previously prepared stocks [1, 6–9].

For cell lysate labelling, cell pellets were resuspended in HR buffer (50 mM Tris-HCl, 5 mM MgCl_2_, 250 mM sucrose, 2 mM TCEP and protease inhibitor t (Roche), pH 7.4), and lysed by sonication (Bioruptor, Diagenode, high intensity for 10 minutes with an ON/OFF cycle of 30 seconds) at 4°C. In SDS-PAGE or immunoblot detection assays, 25-40 μg of clarified cell lysate in 20 μL was labelled with indicated probe (final concentration 1 μM) at 37 °C for 30 min. Reactions were stopped by the addition of LDS (lithium dodecyl sulfate) sample buffer (Invitrogen Life Technologies, Carlsbad, CA, USA) containing 2.5% β-mercaptoethanol, followed by boiling for 7 minutes.

For labeling of purified recombinant enzymes, the enzymes were diluted in assay buffer containing 50 mM Tris-HCl, 100 mM NaCl, 0.5 mg / ml CHAPS and 5 mM TCEP, pH 7.6, to a final concentration of 5 μM. Pure ISG15-PA, and Ub-PA probes were added at a 1:1 molar ratio and incubated for up to 60 min at room temperature. Reactions were stopped by boiling with LDS sample buffer as described above. The recombinant enzymes are listed in Material table 3.

**Material Table 3:**
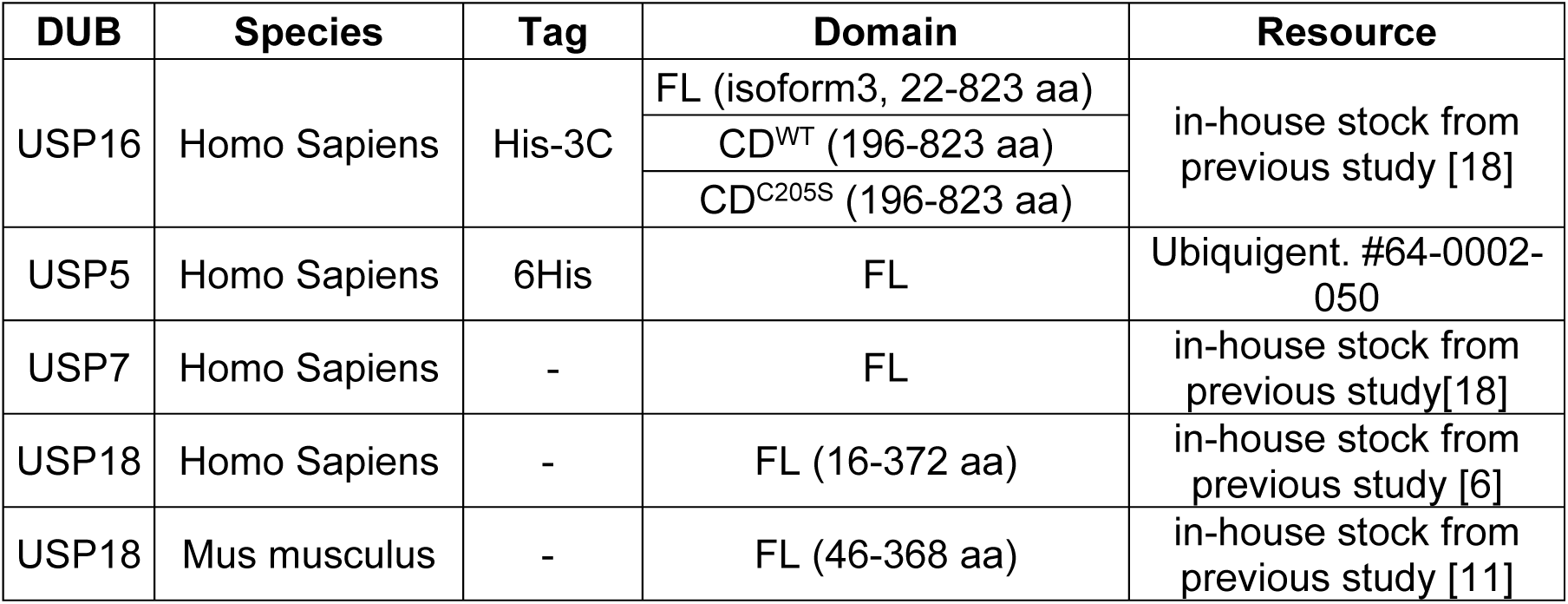
Details of the recombinant enzymes utilised in this study.

| DUB | Species | Tag | Domain | Resource |
| --- | --- | --- | --- | --- |
| USP16 | Homo Sapiens | His-3C | FL (isoform3, 22-823 aa) | in-house stock from previous study [18] |
|  |  |  | CD<sup>WT</sup> (196-823 aa) |  |
|  |  |  | CD<sup>C205S</sup> (196-823 aa) |  |
| USP5 | Homo Sapiens | 6His | FL | Ubiquigent. #64-0002-050 |
| USP7 | Homo Sapiens | - | FL | in-house stock from previous study[18] |
| USP18 | Homo Sapiens | - | FL (16-372 aa) | in-house stock from previous study [6] |
| USP18 | Mus musculus | - | FL (46-368 aa) | in-house stock from previous study [11] |

### DeISGylation assays in cell lysate

HAP1 USP18KO cells or HAP1 USP16KO#B cells were stimulated with 1000 U/ml IFN-β for 48 hours to induce ISGylation. Cell pellet was lysed in EMBO lysis buffer (50mM Tris-HCl pH 7.4, 150 mM NaCl, 2 mM EDTA, 0.5 % NP40, 8mM TCEP) using standard protocols. 40 or 80 μg of cell lysate in 10ul was incubated with recombinant USP16 CD^WT^ or USP16CD^C205S^ at a final concentration of 5uM at 37 °C for 2 hours. The reaction was stopped by boiling with LDS sample buffer as described above.

### ProISG15 cleavage assays

Recombinant human pro-ISG15 protein (R&D Systems, #UL-615-500) was diluted in EMBO lysis buffer (50mM Tris-HCl pH 7.4, 150 mM NaCl, 2 mM EDTA, 0.5 % NP40, 8mM TCEP) to a final concentration of 5 μM, and incubated with 0.5 μM indicated enzymes at 37 °C for indicated time. The reaction was stopped by boiling with LDS sample buffer as described above. The recombinant enzymes are listed in Material table 3.

### ISG15-FP and Ub-FP substrates cleavage assays

The fluorescence polarization (FP) assays were performed as reported before [10, 11]. In brief, two-fold serial dilutions (200–25 nM) of the indicated enzyme solutions (The recombinant enzymes are listed in Material table 3) were prepared in assay buffer containing 50 mM Tris-HCl, pH 7.5, 2 mM DTT, 100 mM NaCl, 1 mg / ml CHAPS and 0.5 mg / ml bovine gamma globulin (BGG). 10 μl of each of the dilution steps was added to the empty wells of the plate (Corning 3820, black, low volume 384 well microplate, LBS, round wells, flat bottom). The reaction was started by addition of 10 μl of the full-length ISG15-FP substrate [10] or the Ub-FP substrate [11] (200 nM final concentration). The fluorescence intensities in the S (parallel) and P (perpendicular) directions were recorded in intervals of 60 or 90 seconds on a BMG Labtech Pherastar plate reader (excitation, 540 nm; emission, 590 nm). From these S and P values, the FP values (in mP) were calculated by adjustment of the FP value (L) of the tracer molecule TAMRA-KG to 50 mP:

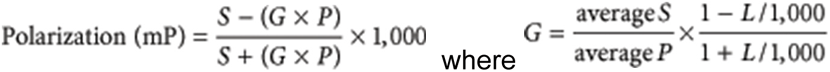

### ISGylation analysis in HEK293T cells

To analyse ISGylation on Myc tagged substrates, ChromoTek Myc trap pulldown assays were performed as GFP trap pulldown before [12]. HEK293T cells, transfected as indicated, were lysed in 300 µL lysis buffer 1 (50 mM Tris-HCl, pH 7.5, 150 mM NaCl, 5 mM ethylenediaminetetraacetic acid (EDTA), 0.5% Triton X-100, 10 mM N-methyl maleimide (general DUB inhibitor diluted in DMSO, freshly added) and protease inhibitors (Roche Diagnostics, EDTA-free, freshly added). Then, 100 µL lysis buffer 2 (100 mM Tris-HCl, pH 8.0, 1 mM EDTA, and 2% SDS) was added to the crude lysates; samples were sonicated (Fisher Scientific FB120 Sonic Dismembrator, 3 pulses, amplitude 40%) and SDS was subsequently diluted by the bringing sample volume to 1 mL with lysis buffer 1. After centrifugation (20 min, 4 °C, 20,817× g), lysates were incubated with 5 µL Myc Trap Agarose (Chromotek) overnight at 4 °C. Beads were washed 5 times with lysis buffer, and denatured with sample buffer by heating at 95 °C for 7 min.

### Electrophoresis and immunoblots

Samples were resolved on precast Bis-Tris NuPAGE Gels (Invitrogen, including 4-12%, 10% and 12% for different samples) using MOPS buffer (Invitrogen Life Technologies, Carlsbad, CA, USA). For fluorescence scan, labeled enzymes were visualized by in-gel fluorescence using a Typhoon FLA 9500 imaging system (GE Healthcare Life Sciences) (Rhodamine channel for probe, Cy5 channel for protein marker). For immunoblots, the gel was then transferred to Nitrocellulose membranes for immunoblots, the signal was visualized using a LICOR Odyssey system. The primary antibodies are listed in Material table 1.

### Activity-based protein profiling (ABPP) with biotin-ISG15-PA probe

Performed as previously described [13]. HAP1 cells were lysed in glass beads lysis buffer (GBL: 50 mM Tris, pH 7.5, 5 mM MgCl2, 0.5 mM EDTA, and 250 mM Sucrose) and protein concentrations determined through a Pierce BCA Assay. Using the results of the BCA Assay, 1 mg of protein from each sample was aliquoted into a clean Eppendorf, and the tubes diluted with glass bead buffer to give V_f_ =∼550 µL, with the need for V_f_ to be the same across all sample tubes. The probe was added to the relevant cell lysates at the pre-determined optimum ratio and all samples incubated: 30 min for the Biotin-ISG15-PA probe (Viva Biotech Ltd) at 37°C. The digest was quenched through the addition of 44 µL 5% (w/v) SDS and 27.5 µL 10% NP-40 to each Eppendorf. Samples were then diluted by adding 1212 µL NP-40 lysis buffer. 150 µL NeutrAvidin Agarose beads (Thermo Fisher) was aliquoted out into one 2 mL Eppendorf tube per sample. The resin was washed with 750 µL NP-40 Lysis Buffer by vortexing briefly and centrifugation (2,000 g, 1 min, RT). The supernatant was then carefully pipetted off and these wash steps repeated four times in total. The samples were added to the resins, sealed with Parafilm and incubated on a turning wheel overnight at 4°C. The following day the samples with resins were spun down (2 000 g, 1 min, RT) and the supernatant carefully removed. The resins were washed with 750 µL NP-40 lysis buffer, vortexed briefly and centrifuged (2,000 g, 1 min, RT). The supernatant was then carefully removed, without disturbing the beads, and these wash steps repeated four times. The probe labelled proteins bound to the agarose beads were eluted by adding 110 µL 2.5x LSLB with 3 mM biotin to the resin. The samples were vortexed briefly and boiled (10 min, 95°C). The beads were pelleted by centrifugation (2 000 g, 1 min, RT) and the supernatant containing the proteins carefully removed without disturbing the resin, and transferred to an Eppendorf tube. Samples are ready for either immunoblotting or LC-MS/MS analysis [13].

For MS analysis, sample eluates were diluted to 175 μL with ultra-pure water and reduced with 5 μL of DTT (200 mM in 0.1 M Tris, pH 7.8) for 30 min at 37 °C. Samples were alkylated with 20 μL of iodoacetamide (100 mM in 0.1 M Tris, pH 7.8) for 15 min at room temperature (protected from light), followed by protein precipitation using a double methanol/chloroform extraction method. Protein samples were treated with 600 μL of methanol, 150 μL of chloroform, and 450 μL of water, followed by vigorous vortexing. Samples were centrifuged at 17,000 × g for 3 min, and the resultant upper aqueous phase was removed. Proteins were pelleted following the addition of 450 μL of methanol and centrifugation at 17,000 × g for 6 min. The supernatant was removed, and the extraction process was repeated. Following the second extraction process, precipitated proteins were re-suspended in 50 μL of 6 M urea and diluted to <1 M urea with 250 μL of 20 mM HEPES (pH 8.0) buffer. Protein digestion was carried out by adding trypsin (from a 1 mg/ml stock in 1 mM HCl) to a ratio 1:100, rocking at 12 rpm and room temperature overnight. Following digestion, samples were acidified to 1% trifluoroacetic acid and desalted on C18 solid-phase extraction cartridges (SEP-PAK plus, Waters), dried, and re-suspended in buffer A.

### ISG15 interactome immunoprecipitation

The whole has been performed as before [1] with the following modifications. HAP1 cells were lysed with Co-IP lysis buffer (20 mM Hepes pH 8.0, 150 mM NaCl, 0.2% NP-40, 10 % glycerol, 5 mM NEM, phosphatase and protease inhibitor cocktails; 25 ×106 cells per condition) and subjected to immunoprecipitation using 5 µg of ISG15 antibody (Boston Biochem #A-380) plus 25 μL of protein G Sepharose slurry (Invitrogen; #15920-10) for 16 h at 4 °C. Beads were washed 4 times with Co-IP lysis buffer and immunocomplexes were eluted with 2X Laemmli. 10% of the eluates was used for immunoblotting with the indicated antibody. The remaining eluate was prepared for MS analysis as previously described (DOI: 10.1126/science.ade8840). Eluted proteins were processed for mass spectrometry analysis using suspension traps (S-Traps). Proteins were reduced with 200 mM DTT in 0.1 M Tris pH 7.8, followed by alkylation with 200 mM iodoacetamide in 0.1 M Tris pH 7.8 in the dark. Samples were acidified by addition of 12% phosphoric acid and captured on S-TrapTM midi columns (C02-midi, ProtiFi). Columns were washed with 90% methanol in 100 mM triethylammonium bicarbonate (TAEB) with centrifugation at 4000 g. Captured proteins were digested with trypsin (1:100 w/w) in 1 mM HCl overnight at RT. Peptides were eluted, dried and dissolved in Buffer A (98 % MilliQ-H20, 2 % CH3CN and 0.1 % TFA).

### Liquid chromatography-tandem mass spectrometry (LC-MS/MS) analysis

LC-MS/MS analysis was performed using a Dionex Ultimate 3000 nano-ultra high-pressure reverse-phase chromatography coupled on-line to a Fusion Lumos (ISG15 interactome) or a Q Exactive (ISG15 ABPP) mass spectrometer (Thermo Scientific) as described previously [13]. In brief, samples were separated on an EASY-Spray PepMap RSLC C18 column (500 mm × 75 μm, 2 μm particle size, Thermo Scientific) over a 60 min (120 min in the case of the matching proteome) gradient of 2–35% acetonitrile in 5% dimethyl sulfoxide (DMSO), 0.1% formic acid at 250 nL/min. MS1 scans were acquired at a resolution of 60,000 at 200 m/z and the top 12 most abundant precursor ions were selected for high collision dissociation (HCD) fragmentation.

### Data analysis

From raw MS files, searches against the UniProtKB human sequence data base (92,954 entries) and label-free quantitation were performed using MaxQuant Software (v1.5.5.1). Search parameters include carbamidomethyl (C) as a fixed modification, oxidation (M) and deamidation (NQ as variable modifications, maximum 2 missed cleavages, matching between runs, and LFQ quantitation was performed using unique peptides. Label-free interaction data analysis was performed using Perseus (v1.6.0.2), and volcano and scatter plots were generated using a t-test with permutation FDR = 0.01 for multiple-test correction and s0 = 0.1 as cut-off parameters.

Other graphs were generated using GraphPad PRISM 8 and Excel and for the statistical analysis, we applied two-way ANOVA tests, including multiple comparison testing via the Dunnett method available through the GraphPad Prism software.

Immunoblot protein bands were quantified using Image Studio Lite Version 5.2.5. quantification Statistical evaluations report on Student’s t test (two-tailed distribution) with *p < 0.05, **p < 0.01, and ***p < 0.001, NS: not significant). All error bars correspond to the mean ± S.E.M.

### Gene ontology and STRING network analysis

The GO consortium web tool (http://geneontology.org/) was utilized to conduct GO analysis. To evaluate enriched GO terms of the identified USP16-dependent ISG15 interactome proteins, the PATHER overrepresentation test (released 20230510) was employed. The proteins were analyzed for overrepresentation of PANTHER GO-Slim biological process, PANTHER GO-Slim cellular component, and PANTHER GO-Slim molecular function terms using the Fischer exact test.

Network analysis of USP16-dependent ISG15 interactome proteins (Data Table S3) was performed using the online STRING database v11.5 (https://string-db.org/). The following setting were applied: Output settings: high confidence interaction score (0.7), edges show protein connections based on textmining, experiments, databases, co-expression, neighborhood, co-occurrence, and gene fusion. The network was exported as a TVS (tab-separated values) file, further analyzed and visualized in Cytoscape version 3.9.1. The Cytoscape plug-in MCODE (molecular complex detection) version 2.0.2 was applied to identify highly connected subclusters of proteins using a degree cutoff of two, cluster finding: haircut, a node score cutoff of 0.2, a K-core of two and a max depth of 100.

## ACKNOWLEDGEMENTS

This work was supported by the Oncode Institute, The Netherlands Organization for Scientific Research (NWO) (VICI grant no. 724.013.002 to H.O.), and the Institute for Chemical Immunology (grant no. ICI00026 to A.S.). The A.P.F. and B.M.K. labs were supported by the Chinese Academy of Medical Sciences (CAMS) Innovation Fund for Medical Science (CIFMS), China (grant number: 2018-I2M-2-002) and by Pfizer.

## AUTHOR CONTRIBUTIONS

A.S., J.G and P.P.G. designed and J.G. performed most of the experiments. A.P.F. and B.M.K. designed experiments involving ABPP and ISG15 interactome analysis (Figure 1, Figure 4 and Supplementary Figures 1, 6 and 7). A.P.F., B.M.K., D.O.B., H.G., and H.C.S. performed sample preparation and analysis for ISG15 ABPP and ISG15 interactome by mass spectrometry analysis and immunoblotting (Figure 1, Figure 4 and Supplementary Figures 1, 6 and 7). D.F. performed experiments in Figure 2A. P.P.G. prepared ISG15 and M20 ABPs, performed experiments in Figure 2B, Supplementary Figure 2. J.J.L.L.A. and J.N. provided plasmids and CRISPR knockout design. K.P.K provided recombinant mouse USP18 protein and anti-human ISG15 serum. G.F. provided recombinant human USP18 protein. J.G., A.P.F., H.v.D., P.P.G. and A.S. interpreted the data and wrote the manuscript, with input from all other authors.

---

**Supplementary Figure 1.**
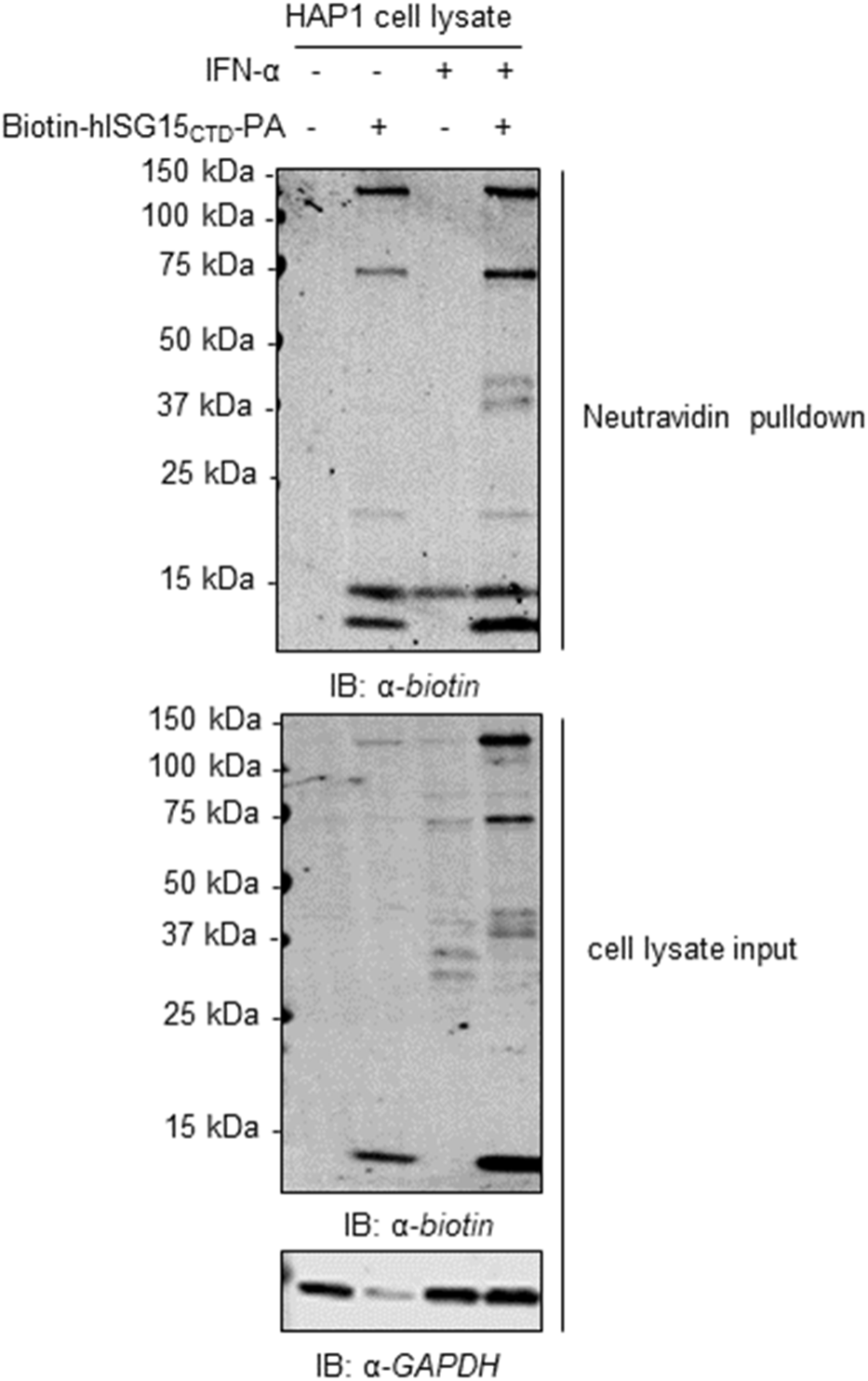
Activity-based probe labelling of ISG15-reactive proteases in human HAP1 WT cells. Anti-biotin immunoblot showing the Biotin-hISG15_CTD_-PA probe enriched ISG15-reactive proteases from human HAP1 WT cells after Streptavidin beads pulldown and denaturing wash steps, corresponding to Figure 1A.

**Supplementary Figure 2.**
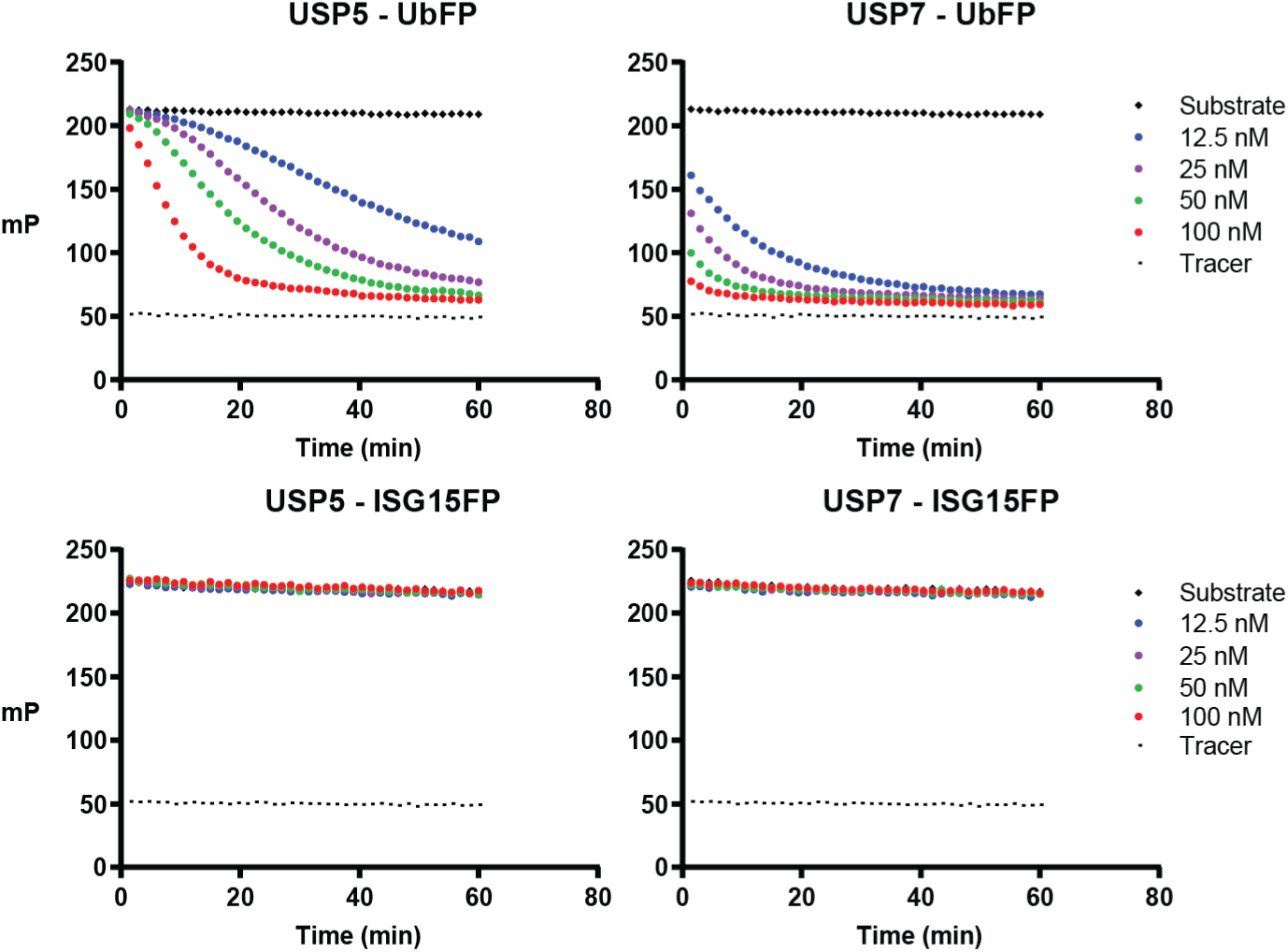
Catalytic activity of recombinant human USP5 and USP7 towards the isopeptide-linked Ub-FP and ISG15-FP substrates. Indicated amounts of USP16 FL/CD enzymes were incubated with 200 nM Ub-FP or ISG15-FP. The substrate cleavage was monitored on the basis of the change in fluorescence polarization (in millipolarization units (mP)).

**Supplementary Figure 3.**
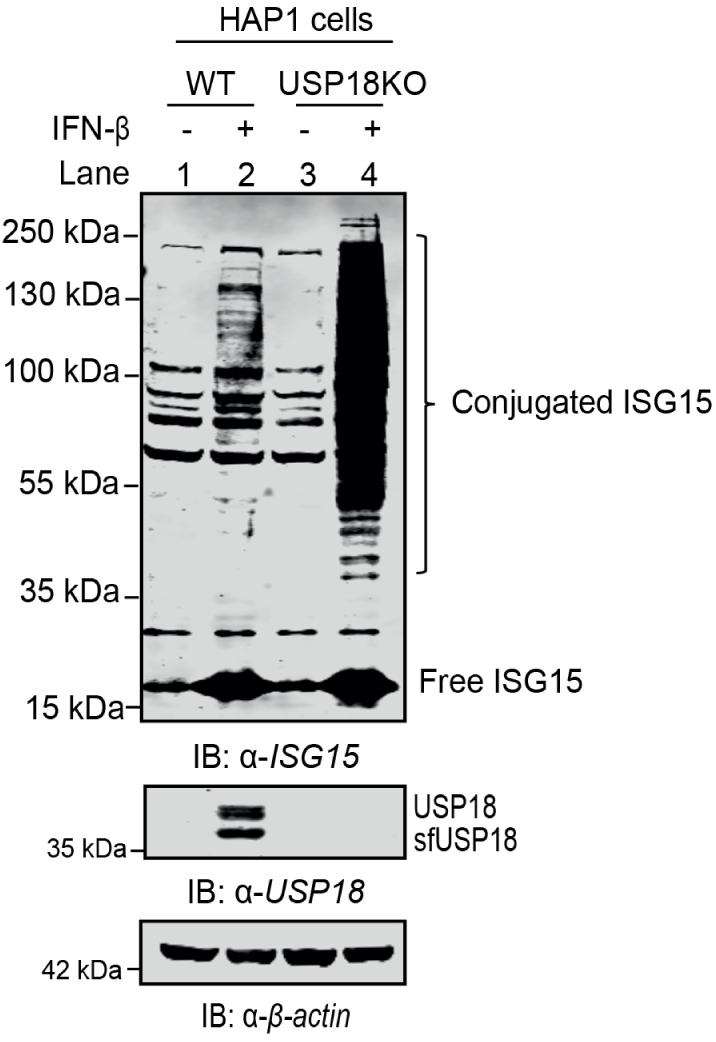
Validation of the HAP1 USP18KO cells. HAP1 WT and USP18KO cells were stimulated with 1000 U/ml of IFN-β for 24 hours to induce ISGylation and USP18 expression. Due to the loss of USP18 in the KO cells, the accumulation of cellular ISGylation was greatly elevated.

**Supplementary Figure 4.**
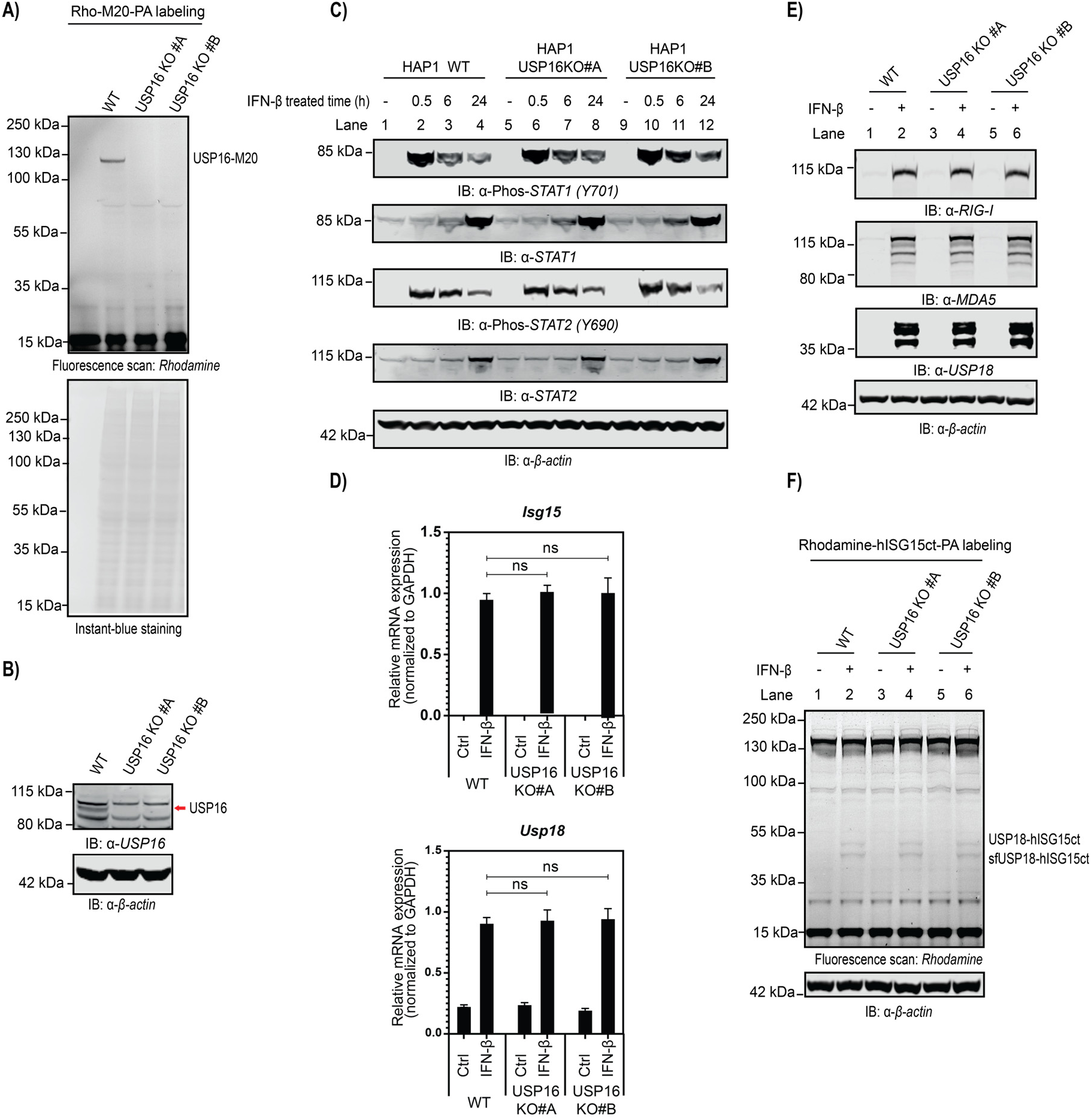
Characterization of USP16 knockout (KO) HAP1 cells generated via CRISPR/Cas9 using guide RNAs targeting exon 6 (KO #A) or exon 4 (KO #B). (A) Endogenous USP16 protein analysis by USP16-specific Rhodamine-M20-PA probe labeling. (B) Endogenous USP16 protein analysis by USP16 immunoblotting in USP16 knockout (KO) HAP1 cells. (C) Immunoblot analysis of type I IFN signaling. HAP1 WT and USP16KO cells were stimulated with 1000 U/ml of IFN-β as indicated, cell lysates were analysed by immunoblotting using the indicated antibodies. (D) Quantitative RT–PCR for *Isg15* and *Usp18* transcripts in HAP1 WT and USP16KO cells stimulated with 1000 U/ml of IFN-β for 24 hours. Bars represent means ± S.E.M with three samples in each group. Data are representative of three independently performed experiments. Immunoblot analysis of selected type I IFN inducible proteins, including RIG-1, MDA5, and USP18. HAP1 WT and USP16KO cells were stimulated with 1000 U/ml of IFN-β for 24 hours, cell lysates were analysed by immunoblotting using the indicated antibodies. ISG15-PA probe labeling of lysates of HAP1 wt and USP16KO cells. Cells were treated with IFN-β (1000 U/ml for 24 hours) or not; lysates were incubated with Rhodamine-hISG15ct-PA probe, resolved with SDS-PAGE, and scanned for Rhodamine fluorescence. Endogenous USP18 is indicated. The β-actin immunoblot confirms equal loading.

**Supplementary Figure 5.**
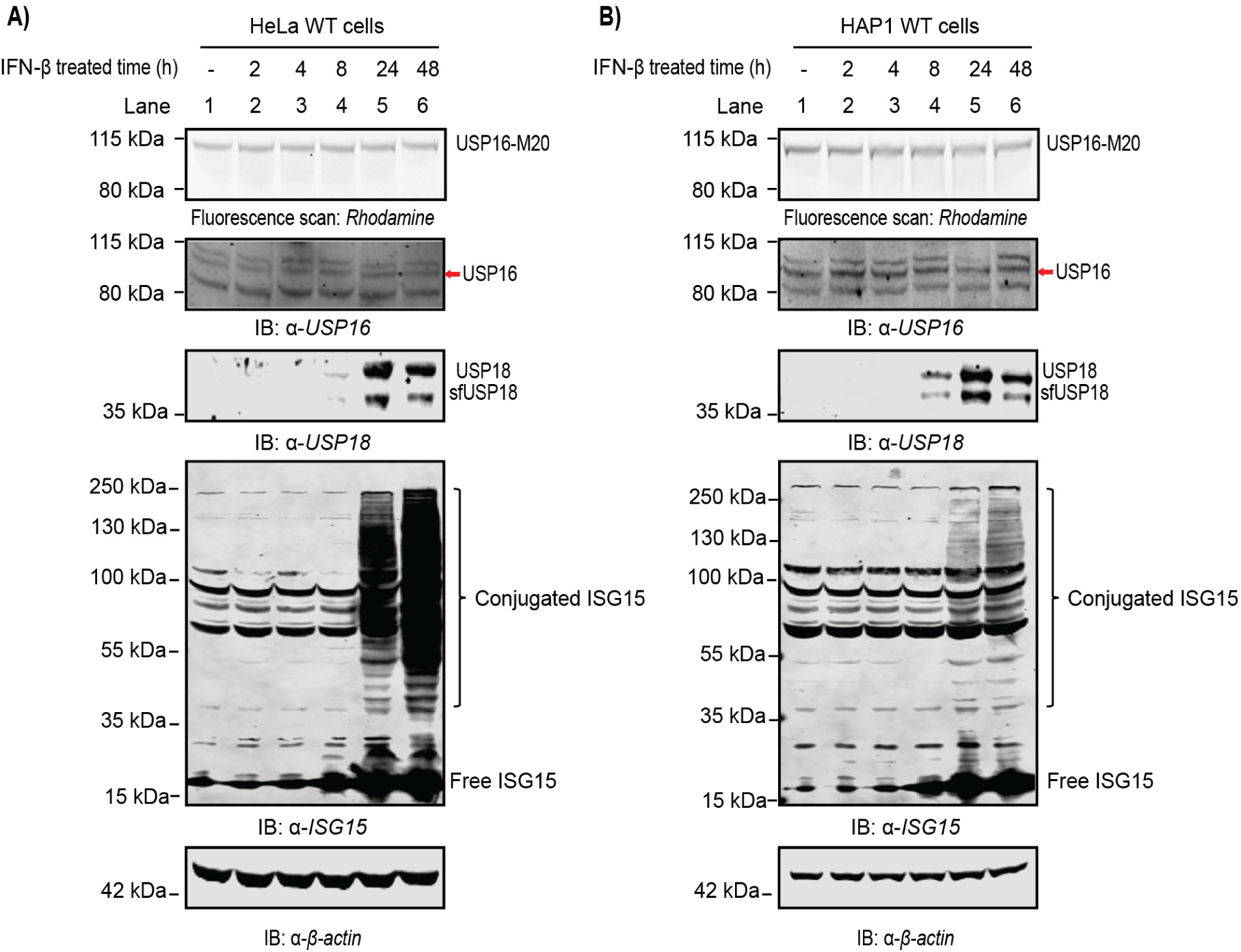
USP16 activity and protein levels are not affected by IFN-β. Assessment of USP16 enzymatic activity and protein expression in HeLa cells (A) and HAP1 cells (B) stimulated with 1000 U/ml of IFN-β as indicated. Cell lysates were labeled by USP16-specific Rhodamine-M20-PA probe, and analyzed by immunoblotting using the indicated antibodies.

**Supplementary Figure 6.**
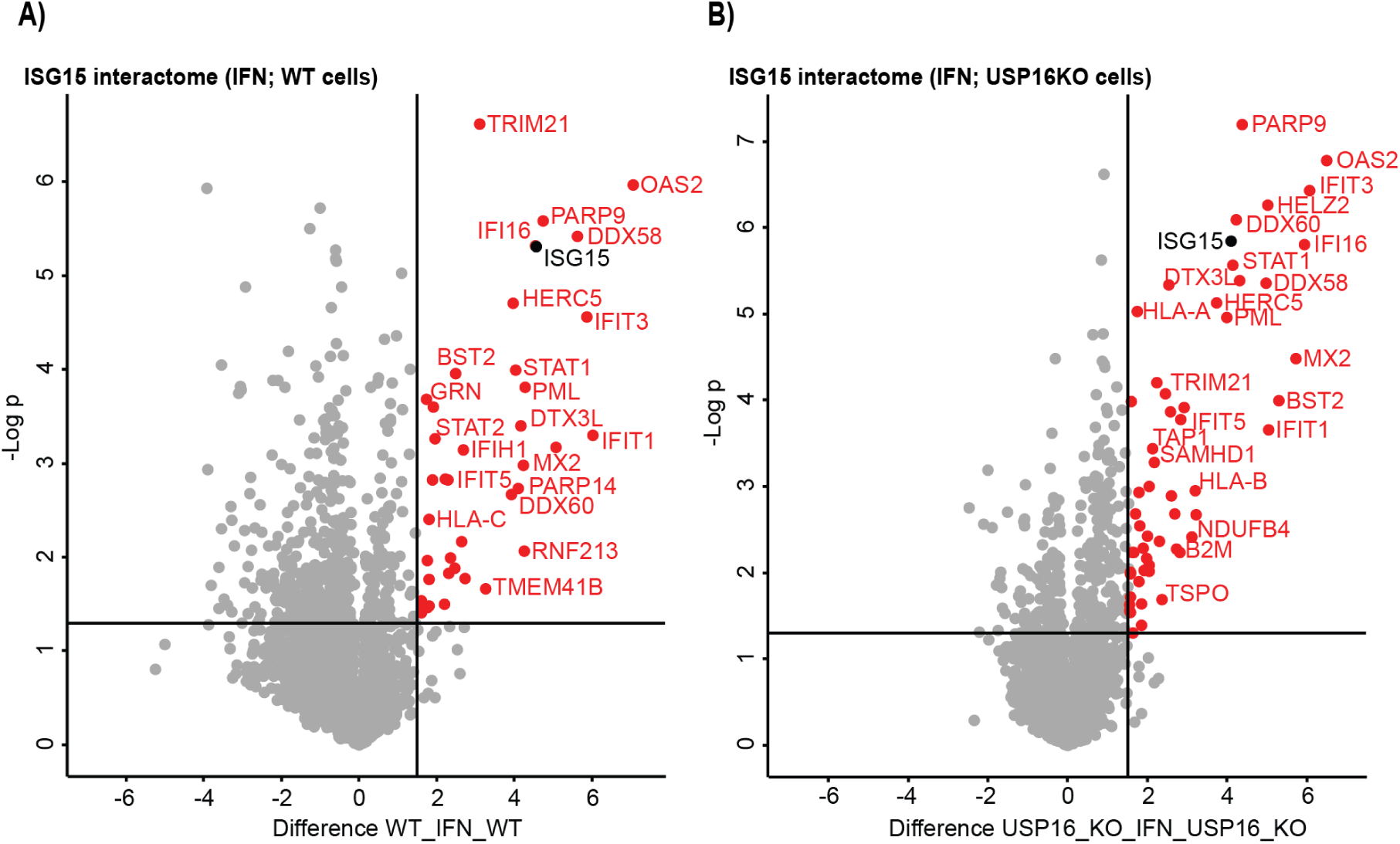
ISG15 interactome in HAP1 WT and USP16KO cells. (A) Volcano plot showing all identified proteins within the IFN-β stimulated WT samples compared with unstimulated WT. Dashed lines indicate a cutoff at twofold change (log2 = 1) and a p-value of 0.05 (−log10 = 1.3), n = 2 independent experiments. Source data are provided as Data Table S4. In red are shown the upregulated proteins in the IFN-β stimulated WT cells, named as “ISG15 interactome in HAP1 WT cells”. (B) Volcano plot showing all identified proteins within the IFN-β stimulated USP16KO samples compared with unstimulated USP16KO. Dashed lines indicate a cutoff at twofold change (log2 = 1) and a p-value of 0.05 (−log10 = 1.3), n = 2 independent experiments. Source data are provided as Data Table S5. In red are shown the upregulated proteins in the IFN-β stimulated USP16KO cells, named as “ISG15 interactome in HAP1 USP16KO cells”.

**Supplementary Figure 7.**
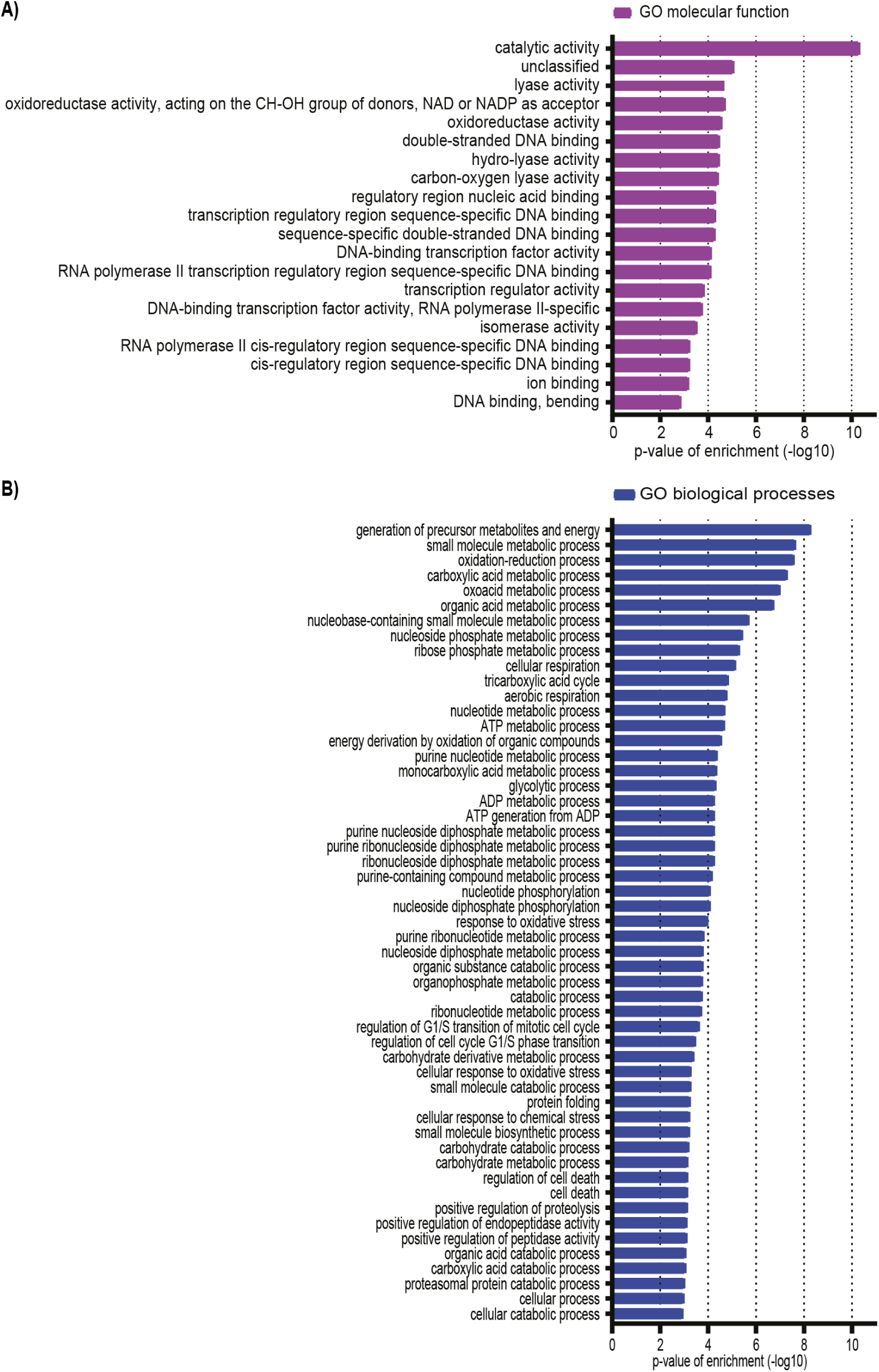
Gene Ontology (GO) enrichment analysis of the USP16-dependent ISG15 interactome with full GO terms. (A) The bar graph shows the full GO terms for molecular functions (MF) of the USP16-dependent ISG15 interactome against the annotated human proteome. (B) The bar graph shows the full GO terms for biological processes (BP) of the USP16-dependent ISG15 interactome against the annotated human proteome.

## Notes

### Competing Interest Statement

The authors have declared no competing interest.

